# LTP-like noninvasive striatal brain stimulation enhances striatal activity and motor skill learning in humans

**DOI:** 10.1101/2022.10.28.514204

**Authors:** Maximilian J. Wessel, Elena Beanato, Traian Popa, Fabienne Windel, Pierre Vassiliadis, Pauline Menoud, Valeriia Beliaeva, Ines R. Violante, Hedjoudje Abderrahmane, Patrycja Dzialecka, Chang-Hyun Park, Pablo Maceira-Elvira, Takuya Morishita, Antonino Cassara, Melanie Steiner, Nir Grossman, Esra Neufeld, Friedhelm C. Hummel

## Abstract

The stimulation of deep brain structures has thus far been possible only with invasive methods. Transcranial electrical temporal interference stimulation (tTIS) is a novel, noninvasive technology that might overcome this limitation. The initial proof-of-concept was obtained through modeling, physics experiments and rodent models. Here, we show for the first time successful noninvasive neuromodulation of the striatum via tTIS in humans using computational modeling, fMRI studies and behavioral evaluations. Theta-burst patterned, LTP-like striatal tTIS increased activity in the striatum and associated motor network. Furthermore, striatal tTIS enhanced motor learning capacity, especially in healthy older participants, who have lower natural learning capacity than younger subjects. These findings suggest exciting methods for noninvasively targeting deep brain structures in humans, thus enhancing our understanding of their functional roles. Moreover, our results lay the groundwork for innovative, noninvasive treatment strategies for brain disorders, in which deep brain structures play key pathophysiological roles.

## Introduction

Neuromodulation of cortical and subcortical brain structures is an important step toward improving our understanding of neuronal processing across brain networks, thereby allowing us to probe and decipher causal brain-behavior relationships^1^. Existing noninvasive brain stimulation (NIBS) techniques, including transcranial magnetic stimulation (TMS) and transcranial electric stimulation (tES), have been widely used to investigate healthy and pathological systems^2^. However, these approaches show a steep depth-focality trade-off^3^, with focality decreasing as the depth increases. As a result, deep brain structures, such as the basal ganglia and hippocampus, cannot be reached directly without diffusely costimulating the overlying cortex^3,4^. Thus, these deep structures have been accessible only through the use of invasive brain stimulation techniques^5^. To perform deep brain stimulation in healthy subjects and reduce the side effects associated with invasive procedures, new concepts and technologies are needed. One exciting possible solution was recently proposed by Grossman *et al*., who introduced the transcranial temporal interference stimulation (tTIS) technique in rodents^6^. During tTIS, two pairs of electrodes are placed on the head, with each pair delivering a high frequency alternating current. Importantly, this frequency should be sufficiently high and thus not affect the mechanisms maintaining neuronal electrical homeostasis. Moreover, a small frequency shift is applied between the two alternating currents. The superposition of the electric fields creates an envelope oscillating at this low-frequency difference, which in turn influences neuronal activity. By optimizing the electrode placement and current intensity ratio across stimulation channels, the maximal amplitude of the envelope can be steered; hence, the primary focus of neuromodulation can be directed toward individual deep brain structures while minimizing neuromodulation in the surrounding and/or overlying areas^6^.

In the present work, we employed the tTIS strategy in humans to study the effects of striatal neuromodulation on local and network brain activity and associated motor learning behavior. Motor learning is a crucial process for a variety of daily life activities ranging from learning to using tools, playing music instruments and recovering from motor disabilities and has been the focus of numerous neuroscientific studies in recent decades^7,8^. These works have revealed that multiple deep brain structures play essential roles in motor learning and motor control, with the striatum playing a central role in this motor network^7,9^. However, in human neuroscience, the roles of these structures have been largely assessed via associative methods, e.g., through indirect inferences from neuroimaging results. In particular, two main motor learning phases have been identified: an initial fast phase, during which subjects significantly increase their performance by integrating sensory inputs, and a later slower phase, during which improvements are less pronounced and gained slower^10^. Moreover, different neural substrates have been suggested to be specifically involved in each of these phases, and the striatum with its substructures has been proposed as one of the most important subcortical hubs that is involved in both the fast and slow learning phases^11,12^. The activation and engagement of different parts of the striatum dynamically change throughout the learning process, with the caudate nucleus implicated during the initial fast learning phase and the putamen more associated with the slow phase^12,13^. Even within the putamen, different compartments have been found to change their activity over time, i.e., the activation shifts from the associative (rostrodorsal) part to the sensorimotor (caudoventral) part during training^14^.

A critical limitation of existing human neuroimaging techniques, e.g., functional magnetic resonance imaging (fMRI), positron emission tomography (PET) and electroencephalography (EEG), is that these approaches provide only associative evidence of the brain-behavior relationships underlying motor learning^1,15^. Most causal evidence originates from animal work^16,17^, striatal lesion studies of patient cohorts^18^, or invasive deep brain stimulation studies of connected nuclei^19,20^, which indicate the significant role of the striatum in motor learning. However, since human data have been obtained from patients with altered network properties due to disorder-related neurodegeneration or lesions, we cannot draw comprehensive conclusions on the physiology of healthy systems. The noninvasive modulation of striatal activity during motor training with the tTIS strategy may allow us to address this critical gap. In the present study, we applied tTIS to the striatum in randomized, double-blind, crossover designs, demonstrating for the first time the possibility of noninvasively targeting the striatum in humans without coactivating overlying cortices beneath the electrodes. Moreover, we characterized local and network effects on brain activity using fMRI recordings during stimulation (Experiment 1) and quantified behavioral effects by studying the evolution and efficiency in acquiring novel hand-based motor skills in healthy young and healthy older subjects (Experiment 2).

## Results

### Experiment 1

#### tTIS increases specific striatal activity during motor learning

In Experiment 1, task-based fMRI was acquired during a sequential finger tapping task (SFTT)^21,22^ with concomitant theta-burst patterned tTIS or high frequency (HF) control stimulation. Theta-burst patterned stimulation was chosen as the active condition because this stimulation has been shown to induce long-term potentiation (LTP)-like effects in previous animal and human works^23,24^. Specifically, a train of two-second theta bursts was delivered every ten seconds to mimic an intermittent theta burst stimulation protocol^24^. The task was divided into six blocks, with each block consisting of ten 30-second repetitions of the sequential finger tapping task with concomitant stimulation alternated with 30 seconds of rest without stimulation.

To investigate the effects of the stimulation on the target region, the average activity in subregions of the striatum was extracted, as shown in Fig. 1a. A significant effect of the region (*F(1, 276)=260.01, p<0.001, pη^2^=0.49* [large]) and a significant region x stimulation interaction (*F(1, 276)=4.48, p=0.035, pη^2^=0.02* [small]) were detected. The region effect can be explained by higher activity in the putamen than in the caudate during the task (*t(276)=-16.13, p<0.001, d=-1.83, Tukey adjustment*). The interaction effect can be explained by higher activity in the putamen during tTIS than during HF control stimulation (*t(276)=-2.55, p=0.01, d=-0.41, Tukey adjustment*), while no difference was observed in the caudate region (*t(276)=0.45, p=0.65, d=0.07, Tukey adjustment*). This result suggests that LTP-like tTIS preferentially increased activity in a striatal subregion (putamen) that was more activated during the motor task. To better understand the effect of stimulation within the putamen, we distinguished the anterior and posterior parts of the putamen (Fig. 1a, bottom panels). The stimulation effect was confirmed, which is consistent with the results reported above (*F(1, 299)=13.47, p<0.001, pη^2^=0.04* [small]), with tTIS leading to increased activity. This increase was not specific to a particular part of the putamen and was present in both subregions as no significant region x stimulation interaction was observed (*F(1, 299)=0.13, p=0.72, pη^2^<0.001* [micro]). Finally, a significant subregion effect was also observed (*F(1, 299)=37.03, p<0.001, pη^2^=0.11* [medium]), with the posterior part of the putamen showing higher activation than the anterior part (*t(299)=-6.09, p<0.001, d=-0.66, Tukey adjustment*).

**Fig. 1:**
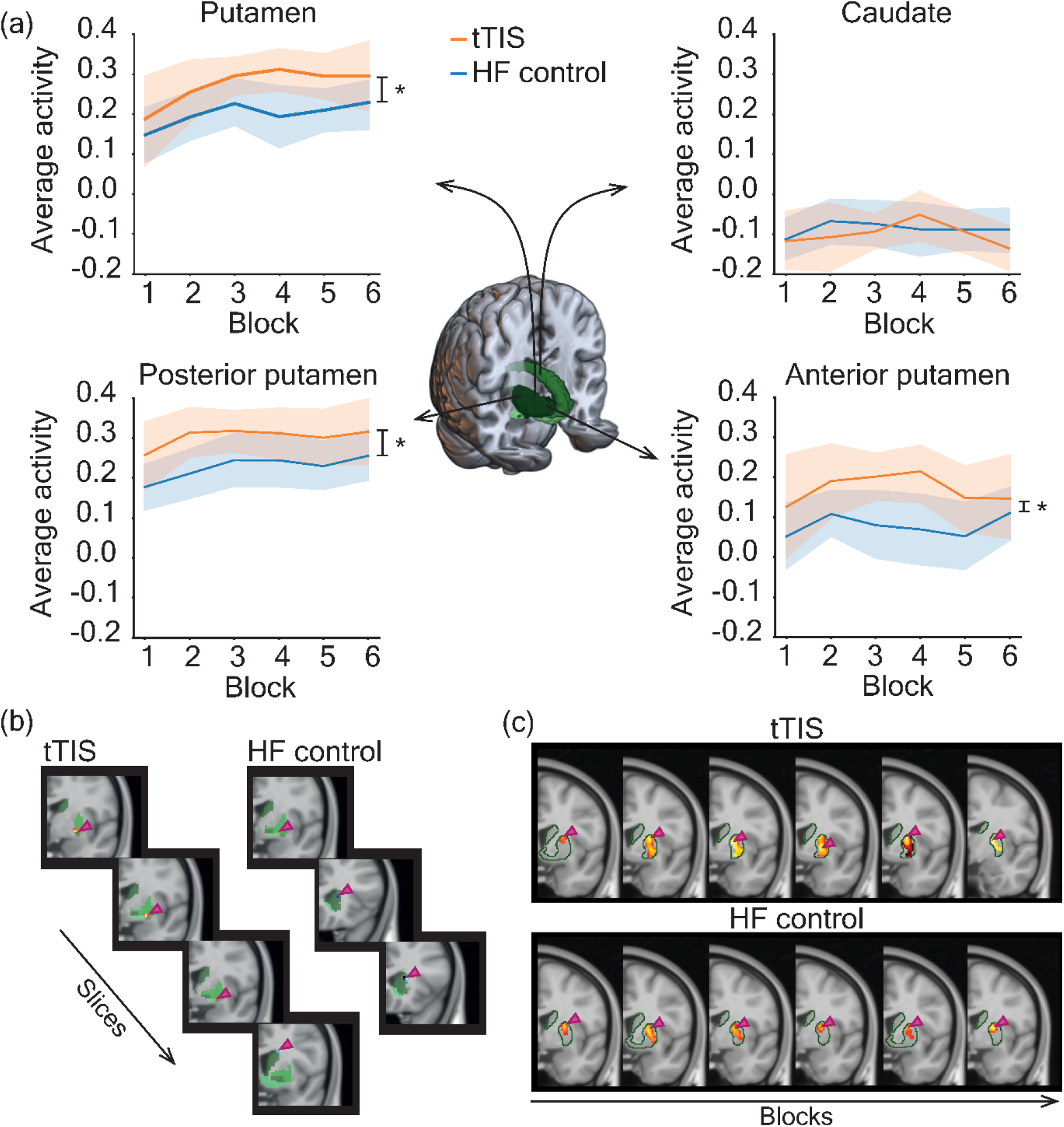
Results of the task-based fMRI experiment – local activity. **(a)** Average BOLD activity in the putamen (top left) and caudate (top right) during tTIS and HF control stimulation. tTIS led to significantly higher activity in the putamen (*t(276)=-2.55, p=0.01, d=-0.41, Tukey adjustment*) but not in the caudate. The average BOLD activity in the posterior (bottom left) and anterior (bottom right) putamen during tTIS and HF control stimulation was also studied. tTIS led to significantly higher activity in both areas (*F(1, 299)=13.47, p<0.001, pη^2^=0.04 [small]*). **(b)** Voxels showing a trend of linear changes (*uncorrected FWE, p=0.01*) over time in the striatum during tTIS are shown on the left and during HF control stimulation on the right. The sections are ordered from caudal to rostral. Hot colors represent increased activity over time, while cold colors represent decreased activity. **(c)** Qualitative characterization of the location of the peak activity during each of the six blocks during tTIS (on top) and HF control stimulation (on the bottom). The left side corresponds to the early training phase, and the right side corresponds to the later training phase. The shift from the superior to the inferior striatum is consistent with previous observations^14,25,26^.

Next, we characterized the temporal changes in activity within the striatum during learning based on previous findings^14,25,26^. Areas in the right striatum showing a trend of linear increases or decreases in activation over time were extracted for each of the two stimulation conditions (*uncorrected voxelwise familywise error (FWE), p=0.01*). Fig. 1b shows that for both stimulation conditions, the activity in the lower part of the putamen increased, while the activity in the superior part of the caudate decreased, which is consistent with the literature^14,25,26^. More in-depth analyses of these functional changes are visualized in Fig. 1c, in which the striatal activity and peak locations are depicted. During tTIS, the activity shifts from the superior striatum to the inferior striatum. This shift was less pronounced when HF control stimulation was applied during the task, with the peak activation still located between the superior and inferior parts of the striatum during the last block of training.

In brief, the analyses indicate that simultaneous application of LTP-like tTIS and motor training induces a differential effect on activity in striatal subregions and accelerates the shift of activation toward sensorimotor subregions, which has been linked with learning in prior studies^14,25,26^. This finding suggests that tTIS can tune learning phase-dependent recruitment patterns in the target region.

#### Striatal tTIS increases activity in the motor network and not in brain regions below the stimulation electrodes

To evaluate how the modulatory effects of striatal LTP-like tTIS influence the rest of the brain, whole brain blood oxygen level-dependent (BOLD) activation during the motor task was compared between the tTIS and HF control stimulation conditions. First, we characterized the regions involved in the motor task during HF control stimulation (Fig. 2a), which included the main nodes of the motor learning network, as expected; for review, please see, e.g., Hardwick and colleagues^7^. Then, clusters with significantly higher activation during tTIS than during HF control stimulation were identified (*uncorrected voxelwise FWE, p=0.001, and corrected cluster-based false discovery rate (FDR)*) (Fig. 2b). Significantly higher activity was found in regions associated with the motor learning network for tasks performed with the left hand, including the right striatum (31.9% of the amygdala cluster), right thalamus and supplementary motor area (SMA), and left cerebellum (for the complete list of regions, please see Table S1 in the supplementary information). In a supplemental analysis, we compared the estimated exposure strength to the tTIS field in supratentorial hubs with higher activity during tTIS to that of the striatal target region. The results indicated higher exposure levels in the striatum, as shown in Fig. 3.

**Fig. 2:**
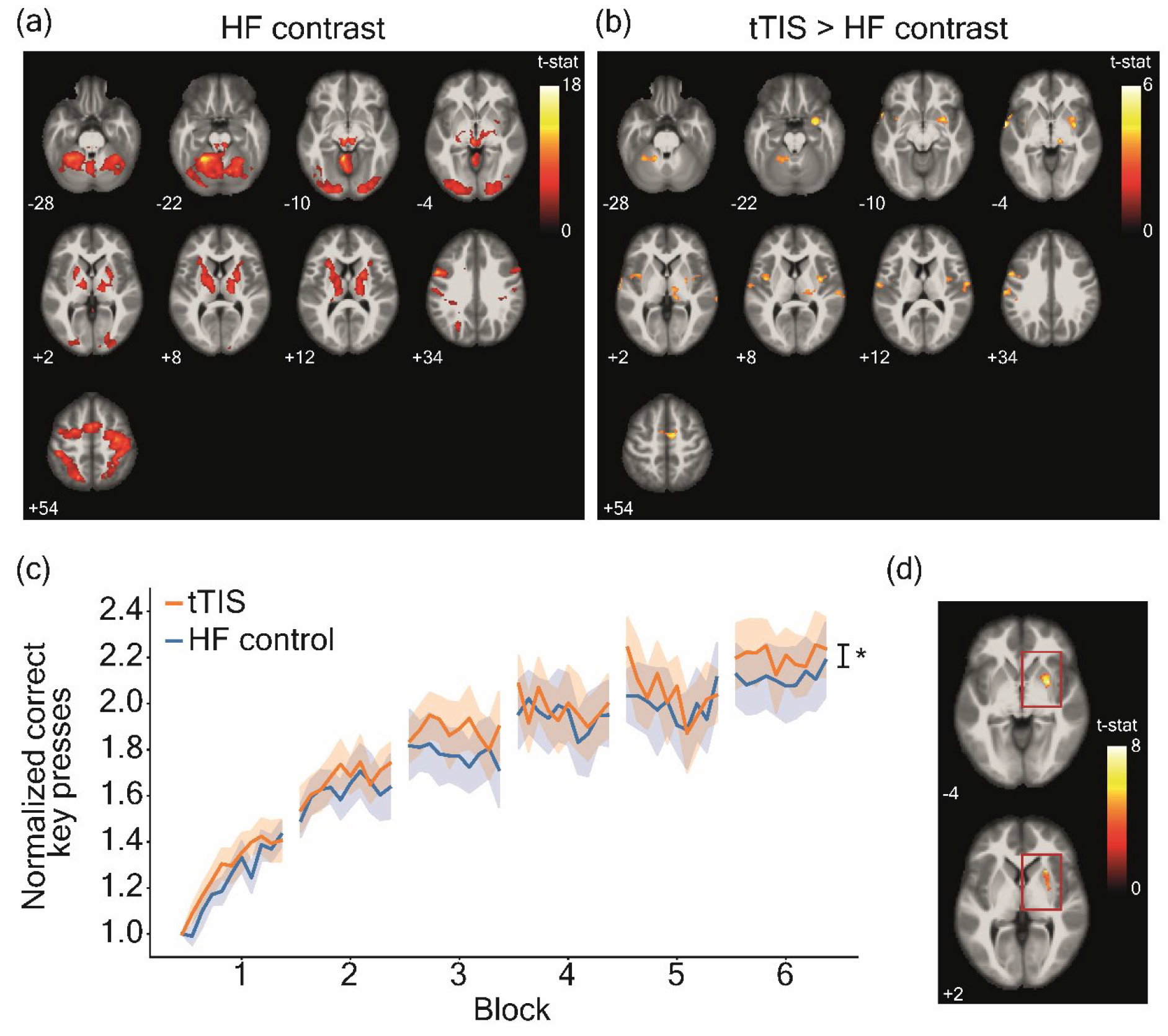
Results of the task-based fMRI experiment – network activity. **(a)** BOLD activity during the motor task with concomitant HF control stimulation. The regions in the motor network involved in the SFTT are shown. Significant clusters are shown for uncorrected voxelwise FWE, *p=0.001*, and corrected cluster-based FDR. **(b)** Comparison of BOLD activity between tTIS and HF control stimulation. Hot colors represent higher activity during tTIS, while cold colors represent lower activity. Significant clusters are shown for uncorrected voxelwise FWE, *p=0.001*, and corrected cluster-based FDR. **(c)** Behavioral results of Experiment 1. The performance is shown as the correct number of key presses normalized to the baseline. A significant effect of the stimulation was present, with tTIS leading to overall higher performance (*F(1, 1560)=6.35, p=0.01, pη2=0.004 [micro]*). **(d)** Areas in the right striatum, where activity was significantly modulated by the behavioral score (correct key presses) during tTIS. Significant clusters are shown for uncorrected voxelwise FWE, *p=0.001*, and corrected cluster-based FDR. No significant clusters were observed during HF control stimulation.

**Fig. 3:**
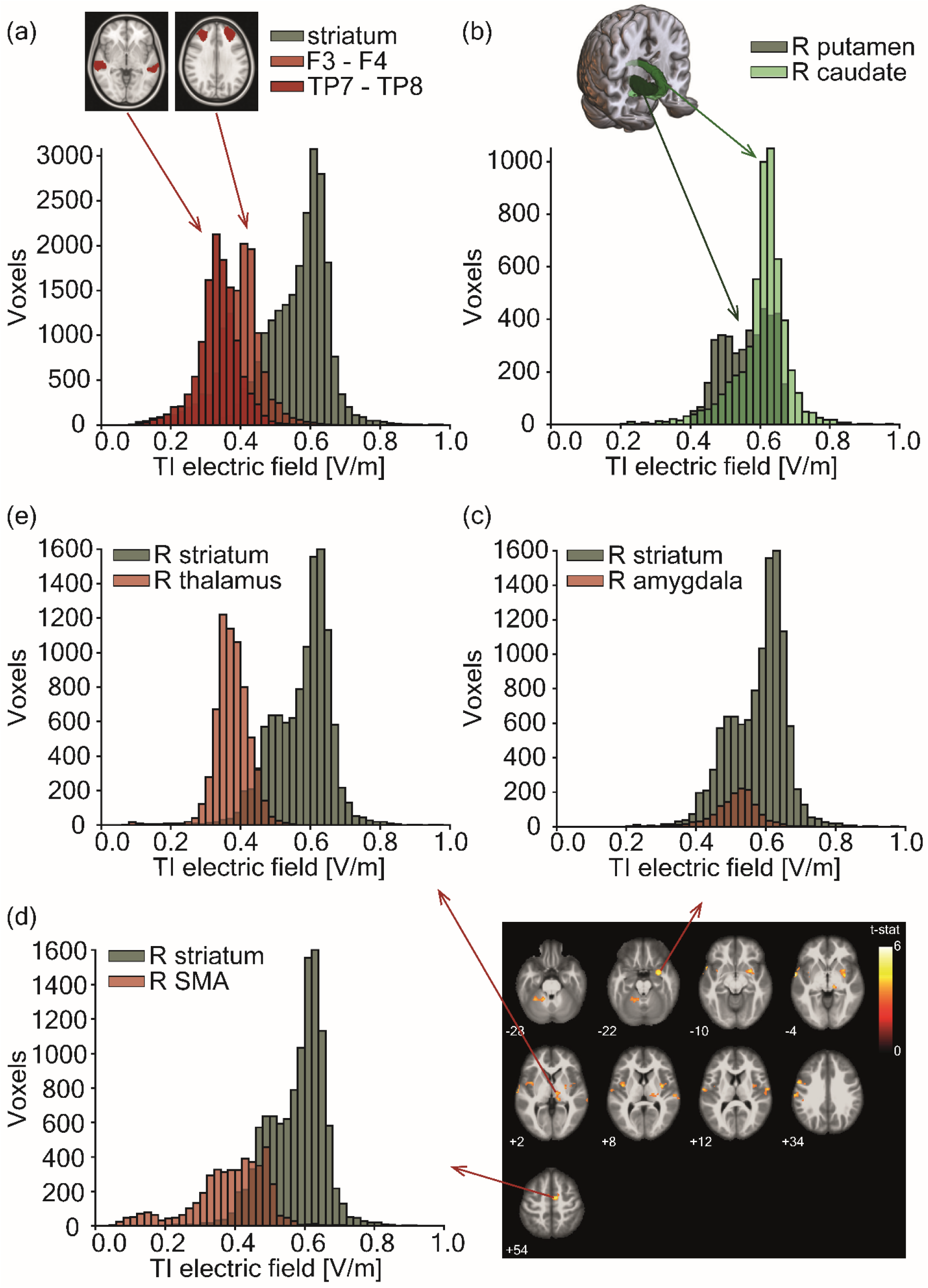
tTIS exposure strength in control regions with respect to the targeted striatum. Histogram depicting the tTIS exposure distribution within specific regions of interest (ROIs) computed for a 2 mA current intensity per channel (peak-to-baseline). **(a)** tTIS exposure distribution of voxels in 10 mm radius spheres underneath the four stimulating electrodes, averaged for the frontal and posterior electrodes, compared to that in the bilateral striatum (putamen, caudate and nucleus accumbens). The horizontal axis scale was limited to the range [0, 1] for visualization purposes. As a result, nine values greater than 1 V/m were omitted, which most likely represented noise values at the edges of the brain mask. **(b)** tTIS exposure distribution of voxels in subparts of the target region, namely, the right putamen and right caudate. (c, d, e) tTIS exposure distribution of voxels in supratentorial hubs showing stronger BOLD activation during the task-based fMRI experiment with concurrent tTIS than during HF control stimulation compared with that in the right striatum (putamen, caudate and nucleus accumbens). **(c)** tTIS exposure distribution of voxels in the right striatum compared to voxels in the specific Brainnetome atlas (BNA^27^) regions of the thalamus, which contained voxels showing higher BOLD activity during the task-based fMRI experiment with concurrent tTIS than during HF control stimulation. **(d)** tTIS exposure distribution of voxels in the right striatum compared to voxels in the specific Brainnetome atlas (BNA^27^) regions of the amygdala, which contained voxels showing higher BOLD activity during the task-based fMRI experiment with concurrent tTIS than during HF control stimulation. **(e)** tTIS exposure distribution of voxels in the right striatum compared to voxels in the specific Brainnetome atlas (BNA^27^) regions of the SMA, which contained voxels showing higher BOLD activity during the task-based fMRI experiment with concurrent tTIS than during HF control stimulation.

In an additional control analysis, we examined whether the BOLD signal below the electrodes was modulated by the stimulation condition. BOLD signals were extracted from 10 mm radius spheres and the region of the Brainnetome atlas (BNA^27^) beneath the electrode location. The following regions in the BNA atlas were selected: the left and right A9/46d (dorsal area 9/46) underlying F3 and F4, respectively, and the left and right anterior superior temporal sulcus (aSTS) underlying TP7 and TP8, respectively. In these control regions, no effect of stimulation was found (*F(1, 651)=2.04, p=0.15, pη^2^=0.003* [micro] when using the sphere model and *F(1, 651)=0.38, p=0.54, pη^2^<0.001* [micro] when investigating activity in the BNA regions); for additional details, please see Fig. S1 in the supplementary information.

These results strongly suggest that striatal tTIS successfully modulated activity in the striatum and the associated motor learning network without engagement of the overlaying cortices beneath the electrodes with respect to the activity during the nonmodulating HF control stimulation.

#### Striatal tTIS increases behavioral responses during motor learning

We next evaluated whether the neural effects of striatal tTIS were associated with changes in motor learning behavior by measuring changes in correct key presses during training (Fig. 2c). Significant effects of block (*F(6, 1560)=243.22, p<0.001, pη^2^=0.48* [large]) and stimulation (*F(1, 1560)=6.35, p=0.01, pη^2^=0.004* [micro]) were found. The significant block factor confirms the presence of learning effects during the task. The small but significant difference between the stimulation conditions highlights that compared with the HF control stimulation, motor task performance improved when tTIS was applied. The block x stimulation interaction was not significant, indicating that stimulation effects did not differ over time. According to these findings, we investigated whether the magnitude of BOLD activation in the striatal target region was related to behavioral outcomes. We considered the normalized number of correct key presses as a parametric modulator in the general linear model at the individual subject level. Group statistics restricted to the right striatum revealed significant modulation of striatal activity in the putamen during tTIS (Fig. 2d and Table S2 in the supplementary material). In contrast, activity was not significantly modulated by behavioral performance when HF control stimulation was applied. This result strengthens the hypothesis of direct modulation of striatal activity via tTIS, which not only leads to higher activation but also supports a relationship between brain activity and behavior.

#### No striatal tTIS effects were observed in the absence of task-evoked activity

Because striatal tTIS was shown to modulate BOLD activity and motor learning, we next assessed whether striatal tTIS modulated BOLD signals in the absence of task-evoked activity. Resting state fMRI (rs-fMRI) was acquired in separate sessions. We analyzed seed-based connectivity using the right striatum as a seed. We did not observe a difference in connectivity between striatal tTIS and HF control stimulation (*uncorrected voxelwise FWE, p=0.001, and corrected cluster-based FDR*). The absence of significant effects supports the hypothesis that behavioral coactivation is necessary to induce LTP-like theta burst tTIS effects in the targeted brain area, linked network and associated behavior. This is in line with what has previously been suggested for other conventional low-intensity plasticity-modulating tES protocols^28^.

### Experiment 2

#### Striatal tTIS effects are larger in older adults

In a second experiment, we validated the striatal LTP-like tTIS approach in a behavioral experiment by recruiting a cohort of older adults (N=15, right-handed, 9 females, average age 66.00±4.61 years), who often demonstrate diminished performance gains in motor learning tasks^29,30^, have less tuned underlying brain networks^31,32^ and may have higher sensitivity to plasticity-modulating NIBS protocols^29,33^. Additionally, a new cohort of young healthy control subjects (N=15, right-handed, 8 females, average age 26.67±4.27 years) was recruited.

The subjects performed the SFTT in a shorter training session with longer blocks (seven 90-second blocks) while simultaneously receiving either striatal tTIS or HF control stimulation following a randomized, double-blind, crossover design. The overall duration of the training and stimulation was approximately three times shorter than that in Experiment 1. This difference was introduced to adapt the protocol for follow-up studies recruiting patient cohorts and to homogenize the protocol with protocols considered in previous studies^22,29^.

The analysis of the training phase indicated significant effects of the block (*F(6, 351)=30.16, p<0.001, pη^2^=0.34* [large]) and population (*F(1, 27)=4.36, p=0.046, pη^2^=0.14* [large]), as well as significant stimulation x population (*F(1, 351)=6.71, p=0.01, pη^2^=0.02* [small]) and block x population interaction effects (*F(6,351)=2.29, p=0.04, pη^2^=0.04* [small]) (Fig. 4a and b). Post hoc analysis of the stimulation x population interaction indicated a significant difference across stimulation conditions in the older cohort, with this cohort performing better during striatal tTIS than during HF control stimulation (*t(351)=3.26, p=0.001, d=0.45, Tukey adjustment*). No significant difference was found in the younger cohort (*t(351)=-0.45, p=0.65, d=-0.06, Tukey adjustment*). Moreover, we investigated behavioral changes by computing the gain as the difference between the first and last task blocks. A significant difference was found for the older cohort, with striatal tTIS leading to significantly higher gains than HF control stimulation (V*=96, p=0.04, d=0.76*). No significant difference was found for the younger cohort (*V=40, p=0.28, d=-0.39*). The stimulation-associated effect in the younger and older cohorts was specific to the trained motor sequence, as no effects on motor performance were detected in an intermingled block, in which the order of key presses followed a predefined pseudorandom sequence (younger subjects: *V=53, p=0.72, d=-0.19;* older subjects: *V=82, p=0.23, d=0.44);* for additional details, please see Fig. S2e and f in the supplementary information.

**Fig. 4:**
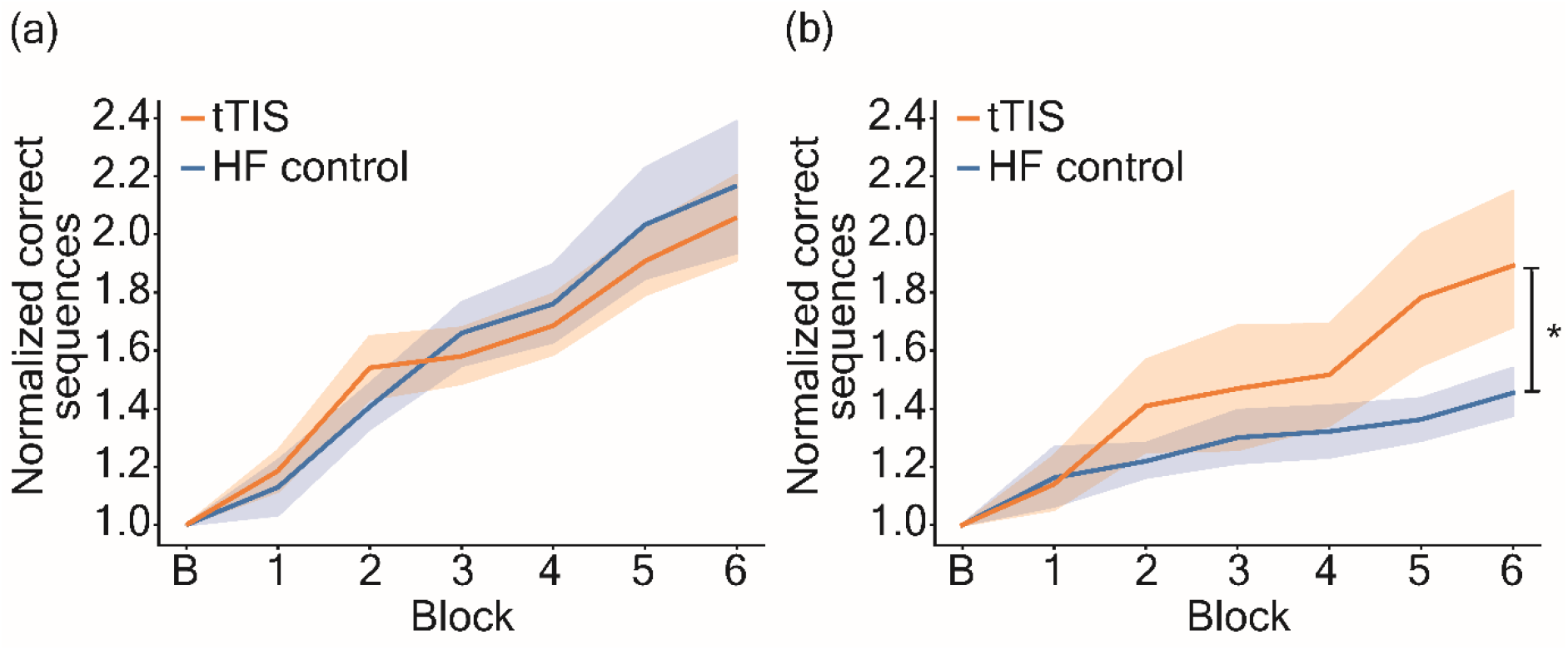
Behavioral experiment results. **(a)** Motor task performance of the younger cohort. The performance is shown as the correct number of sequences normalized to the baseline. No differences across stimulation conditions were observed. **(b)** Motor task performance of the older cohort. The performance is shown as the correct number of sequences normalized to the baseline. The post hoc analysis showed that this cohort performed significantly better during tTIS than during HF control stimulation (*t(351)=3.26, p=0.001, d=0.45, Tukey adjustment*).

In both cohorts, the effect of stimulation on the follow-up measurements (post training (post), 90-min follow-up (FU1) and 24 hours follow-up (FU2)) was also investigated, by analyzing performance normalized to the last block of the training. No significant stimulation effect was found (stimulation factor: *F(1, 537)=0.15, p=0.70, pη^2^<0.001* [micro]; stimulation x follow-up: *F(3, 537)=0.22, p=0.88, pη^2^=0.001* [micro]; stimulation x population: *F(1, 537)=2.71, p=0.10, pη^2^=0.005* [micro]).

#### Analyses of stimulation-associated sensations, blinding integrity, attention and fatigue

The analyses of stimulation-associated sensations across stimulation conditions did not reveal differences in strength across the tested current intensity levels (0.5, 1.0, 1.5, and 2 mA per stimulation channel) (stimulation condition x current strength interaction, *F(3,837.95)=0.06, p=0.98, pη^2^<0.001* [micro]) or sensation categories (stimulation condition x sensation category, *F(6,727.26)=0.73, p=0.63, pη^2^=0.006* [micro]); for additional details, please see Fig. S3 in the supplementary information. At the end of the experiment, the subjects correctly identified the session in which tTIS was applied at approximately chance level (Experiment 1: task-based fMRI *p=0.75*, rs-fMRI *p=0.55;* Experiment 2: *p=1.00* for both cohorts); for additional details, please see the supplementary information. This finding suggests the excellent blinding integrity of tTIS. Furthermore, the stimulation and time did not alter the subjects’ attention (Experiment 1: *V=45, p=0.66;* Experiment 2 – young: *t(14)=-0.54, p=0.60;* Experiment 2 – older: *V=35, p=0.48*) or fatigue levels (Experiment 1: *t(13)=-0.77, p=0.46;* Experiment 2 – young: *t(14)=-0.77, p=0.46;* Experiment 2 – older: *t(14)=-1.55, p=0.14*), as quantified with visual analog scales; please see Fig. S4 in the supplementary information for more information.

## Discussion

The present study demonstrates for the first time that LTP-like striatal tTIS can noninvasively modulate striatal activity and improve motor learning in humans by investigating three independent cohorts (two younger cohorts and one older cohort). Specifically, striatal tTIS enhanced activity in the putamen and in core hubs of the associated brain network. Furthermore, striatal tTIS led to behavioral effects by increasing training gains during a motor learning task. The behavioral effect was particularly pronounced in older participants, who have lower motor learning capacity^29,30^ and less well-tuned underlying brain networks than younger participants^31,32^. In this work, we demonstrate that tTIS can overcome the depth-focality tradeoff observed in conventional NIBS techniques in humans, leading to sufficient current strengths to induce specific, focal and functionally relevant modulation of brain activity in deep brain structures such as the striatum.

An important feature of tTIS is that it operates in the subthreshold range and does not directly induce neuronal action potentials. Thus, to further shape its topographic specificity, behavioral coactivation is likely needed. This argument is supported by the finding that striatal tTIS did not modulate seed-based functional connectivity in the target region during resting state, which was quantified by concurrent rs-fMRI recordings. In other words, when the target region is at rest, tTIS alone cannot affect its connectivity. Furthermore, whole-brain analyses of task-evoked fMRI activity indicated tTIS-associated increases in functional activation only in the right striatum (Fig. 2b), which is strongly engaged in motor learning paradigms performed by the contralateral left hand^14,34^. This result is consistent with previous theories and experimental data acquired with brain slice-based electrophysiology, which suggests that the generation of neuroplasticity through low-intensity tES protocols requires coactivation by synaptic input or task-induced activity^35,36^. In addition, the presence of endogenous activity has been shown to lower the entrainment threshold of neuronal activity at certain resonant frequencies^37,38^.

How are the brain activation patterns induced by striatal LTP-like tTIS linked to the associated behavioral enhancements? The present data suggest multiple, potentially complementary phenomena. First, the correlation analysis in the right striatum suggests that the magnitude of the tTIS-induced BOLD activity in the right putamen is associated with behavioral performance (Fig. 2d). Thus, stronger tTIS-associated activation in this specific target subregion is beneficial for supporting motor learning behavior. Second, the region of interest (ROI)-based analysis indicates that tTIS increased activity in the putamen, which comprises a large part of the sensorimotor subdomain of the striatum (Fig. 1a)^39^, which has been linked to a more advanced stage of motor learning^14,25,26^. These stimulation-induced effects on activity were not observed in the caudate nucleus, which is linked to the associative subdomain (Fig. 1a)^39^. Third, striatal tTIS facilitated the learning-phase-dependent activity shift toward inferior sensorimotor subregions over time (Fig. 1b and c). Overall, these findings suggest that striatal tTIS increases activity in striatal subregions linked to advanced learning phases^14,25,26^, thereby enhancing associated behaviors.

Does striatal tTIS achieve focused neuromodulation with minimal exposure in the overlying cortices or other functionally relevant hubs of the motor learning network? The present results highlight stronger activation in typical core areas of the task-related motor network^7^ during tTIS. Even though this could be due to off-target stimulation, the likelihood is low based on the tTIS field modeling results, which indicate lower exposure in these hubs than in the striatum, as shown in Fig. 3. Moreover, we note that similar results could have been found by stimulating cortical areas. However, it is again highly unlikely that the lower tTIS fields exposure observed in the cortical areas, namely the BNA regions beneath the electrodes, could induce the observed activity difference (please see Figs. 3 and 5f). This result was further confirmed by dedicated control analyses, which indicated that striatal tTIS does not modulate BOLD signals in BNA regions below the electrodes (please see Fig. S1). Finally, the other possible component of the stimulation, namely, the HF control fields, was comparable in the tTIS and HF control stimulation; thus, this component should lead to similar effects in the brain in both conditions, which is not consistent with the differences in the observed BOLD signal. Our findings support a causal relationship between tTIS-induced changes in striatal activity and the connected areas of the motor network, in which activity was enhanced as a function of the striatal stimulation in the absence of significant stimulation in the overlaying brain regions.

**Fig. 5:**
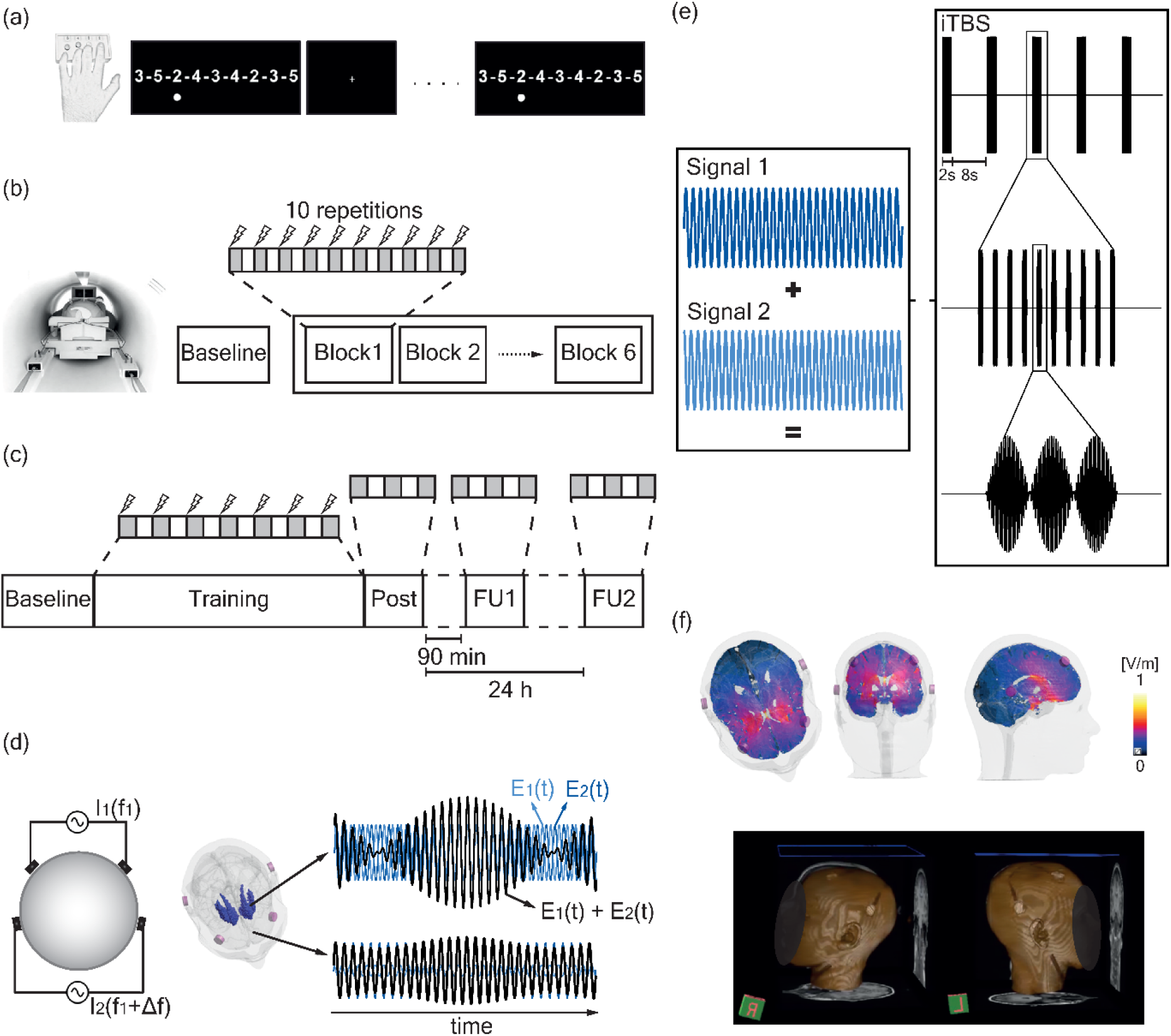
Experimental setup. **(a)** Illustration of the sequential finger tapping task (SFTT)^21,22^: participants were asked to reproduce a sequence displayed on a screen with a consisting of four buttons box. **(b)** Protocol of the task sessions in Experiment 1: the baseline measurement was followed by training with concomitant stimulation, consisting of 6 blocks containing 10 30-second task repetitions alternated with 30-second rest. **(c)** Protocol of Experiment 2: the baseline measurement was followed by training with concomitant stimulation, consisting of 7 repetitions of 1 minute 30 second task blocks. A post assessment with 3 block repetitions was performed immediately after, ~ 90 minutes after and ~ 24 hours after the end of the stimulation. **(d)** Temporal interference stimulation concept. On the left, two pairs of electrodes are shown on a head model, and currents are applied with frequencies of *f_1_* and *f_1_+Δf* On the right, the interference between the two electric fields within the brain is plotted at two different locations with high and low envelope modulation. **(e)** Intermittent theta-burst pattern tTIS is obtained by combining the two high-frequency currents (trains of 2 seconds, repeated every 10 seconds, composed of 10 bursts at 5 Hz, with each including 3 pulses at 100 Hz). (f) Electric field modeling with the striatal montage. The top panel shows the electric tTIS exposure distribution in three chosen slices passing through the target region. The bottom panel shows a 3D reconstruction of the structural MRI images, highlighting the electrode positioning.

Does the tTIS effect depend on the stimulation dose or the lifespan stage? The present findings indicate a dose-dependent effect of striatal tTIS stimulation on behavior. In younger subjects, motor performance increased when the stimulation was applied for up to half an hour (in Experiment 1), which is three times the dosage applied in Experiment 2, during which no stimulation effects were observed. This difference could be due to an already optimal integration of task-relevant information, which is mainly important during early learning stages. The stimulation may thus support the optimization process of motor sequences during later stages of online learning via the cortico-basal ganglia loop^40^. Despite the shorter protocol, when investigating the striatal tTIS effects in an older population, the motor performance during training was clearly better during tTIS than during HF control stimulation. Striatal LTP-like tTIS led to strongly enhanced learning effects, with a 33.6% improvement over the control condition, even with this short training protocol. The pronounced response to the present intervention in the older participants can be explained by several potential reasons. One simple explanation is that healthy older adults show more room for improvement with striatal stimulation than young adults since healthy older adults show decreased performance levels and motor learning abilities^29,30^. Another possible explanation is based on previous imaging studies, which suggest suboptimal processing across dedicated brain networks during motor learning in older adults^41,42^. Thus, striatal tTIS might improve processing in this striato-cortical network and lead to corresponding behavioral improvements. Moreover, aging is related to structural and functional neurodegeneration and reduced brain plasticity, which are in turn associated with functional impairment^43,44^. Improving brain plasticity in regions affected by aging-related changes might result in restoring “natural” dynamics, ultimately leading to behavioral improvements^29^. In line with this hypothesis, previous studies on neurological disorders also found stronger NIBS effects in patients showing stronger dysfunction and impairment^45^. Thus, the observed results suggest that striatal LTP-like tTIS might have larger behavioral effects in cohorts with more pronounced brain malfunctions, such as in healthy older individuals and patients with brain lesions or neurodegenerative disorders. However, this hypothesis has not yet been tested.

What are the possible underlying mechanisms of striatal tTIS? In recent decades, several studies have consistently demonstrated that LTP-like protocols can induce effects on brain plasticity^46–48^. Early evidence of modulatory effects originated from work conducted in brain slices, in which LTP was observed when two high-frequency bursts were applied with interpulse intervals between 200 ms and 2 seconds^23^. The first burst was hypothesized to act as a primer of postsynaptic activity. Hence, by manipulating stimuli timing, either LTP or long-term depression (LTD) can be induced based on the postsynaptic state. The invention of patterned TMS enabled LTP-/LTD-like protocols to be applied in in vivo studies on monkey and human subjects. In this case, theta-burst patterned protocols showed the ability to modulate cortical brain activity and plasticity in an LTP-/LTD-like manner^24,49^. Here, we applied tTIS to achieve comparable LTP-like effects in the striatum. After the introduction of the temporal interference concept in the brain stimulation field by Grossman *et al*.,^6^ the findings were reproduced in animal and computational models^50–52^, however, further investigations of the underlying mechanisms led to several hypotheses, open questions, and disagreements between mechanistic models and experimentally observed responses^53^. For instance, experimental findings suggest that the stimulation effects depend on time constants of axons’ membrane and slow GABAergic inhibition (GABAb-type)^53^. indicating selective responsiveness depending on neuron types and properties^54^.

GABAergic receptors, including GABAb-type, are highly expressed in the striatum, with evidence pointing toward expression of GABAb receptors on dopaminergic neurons^55^, which are important for the occurrence of striatal LTP-effects^56^. Although LTP/LTD-like plasticity effects have been mainly studied in hippocampal and cortical slices, there is strong evidence that comparable phenomena occur in the basal ganglia. Previous work has shown that theta-burst patterned stimulation can induce LTP- and LTD-like effects in the striatal dominant cell type, the GABAergic projecting medium spiny neurons (MSNs), especially in the cortical-striatal and thalamo-striatal inputs^57^. The results of the present study can thus be interpreted as demonstrating the ability of striatal tTIS to induce LTP-like effects and optimize the integration of cortical and thalamic inputs in the striatum, thereby enhancing the effect of excitatory thalamo-cortical projections. This hypothesis is further supported by the present imaging data, which show enhanced activity in the striato-cortical motor network during tTIS. Thus, the current effects are most likely implemented in the direct-pathway circuit, whose net effect would lead to a disinhibition of excitatory thalamo-cortical projections, which would in turn activate cortical premotor/motor circuits^58^. Moreover, an additional characteristic of the axonal membrane, which is fundamental in tTIS, is the passive membrane filter property of neurons^59^. Currently, there is an open debate with alternative mechanistic hypotheses affirming the necessity of a rectification step prior to filtering to demodulate the electric field, thus allowing selective responses to the modulating envelope^60^. To date, only a small number of published studies have applied tTIS in humans^61–64^. Still, they have targeted primary motor and parieto-occipital cortices, which are already reachable with conventional NIBS techniques. The studies targeting motor cortex suggest that tTIS can obtain neural and behavioral effects, namely, tTIS can modulate functional resting-state connectivity, induce faster reaction times and increase implicit motor learning^61,62^. Furthermore, Violante et al. (*preprint 2022, bioRxiv*) suggested that theta-band tTIS applied to the hippocampus can modulate the activity and the connectivity profile of the subcortical target structure and enhance episodic memory performance in young healthy subjects^65^. Importantly, electric field modeling and measurements in a human cadaver suggest that this effect is driven by focused stimulation of the hippocampus and minimizes tTIS exposure in the overlaying cortex.

Together with our results this suggests that tTIS can focus on specific deep brain regions in human subjects without engaging overlaying cortices. These effects are induced by the temporal interference modulation and are independent of the HF content of the carrier signal. tTIS can modulate brain activity in the target region, the associated brain network and linked behavior. Importantly, striatal tTIS induces only minimal stimulation-associated sensations, has good blinding integrity and does not modulate subjects’ attention or fatigue levels (please see the supplementary information), which is an important prerequisite in future controlled human neuroscience and clinical studies.

In addition to the current findings, several points should be addressed. First, even though the present intervention was applied in three cohorts, including both younger and older individuals, and across two experiments, interpretations of these findings need to consider the small sample size. Second, NIBS techniques show relevant intersubject variability in terms of response rates^66^. The degree of the stimulation response variability during tTIS within and across subjects is currently unknown and should be addressed in future studies. Third, based on work suggesting mild effects of low-intensity kHz-frequency stimulation at the cortical level, we cannot rule out the possibility that a portion of the induced striatal neuromodulation effects was caused by the unmodulated high-frequency signal^67^. However, the fact that the montage for the striatal target was optimized based on the modulated tTIS fields and that we could detect several significant contrasts between the tTIS and the HF control stimulation strongly suggests a decisive contribution of the time-modulated exposure magnitude. Finally, in the present work, the optimized electrode montage was chosen on the basis of electric field distributions from simulations involving a detailed reference head model. However, there are important variabilities in anatomy and tissue properties, which can explain some of the observed response variability^68^. By using image-based and subject/patient-specific modeling to personalize electrode placement and stimulation parameters, it is likely that the selectivity and effectivity of tTIS could be further optimized^69,70^. Subject-specific, image-based information about brain anisotropy, e.g., from DTI, can also be used to consider the known^68,71^ impact of the relative orientation of the exposing field and the principal neural structures on stimulability^68^, in addition to the stimulability differences inherent to the different brain regions and neuron types.

## Conclusion

The present work reveals for the first time in humans the ability to noninvasively modulate neuronal activity in deep brain regions via theta-burst patterned LTP-like striatal tTIS. The modulation led to increased activity not only in the targeted deep brain structure, namely, the striatum, but also in the linked functional brain network. Furthermore, striatal LTP-like tTIS induced significant behavioral improvements in a motor learning task, and this effect was especially pronounced in healthy older subjects.

In general, the proposed stimulation approach is a crucial step forward for the field of systems neuroscience as it allows to characterize effects of direct neuromodulation of deep brain activity noninvasively. This approach thus suggests exciting opportunities for better understanding physiological and pathophysiological processes based on causal rather than associative evidence, e.g., evidence derived using conventional neuroimaging techniques.

Overall, the proposed tTIS approach has high potential in noninvasively modulating and studying brain plasticity of deep brain structures in clinical contexts. This is of particular interest and importance as deep brain regions, such as the striatum, hippocampus and thalamus, play critical roles in various motor and cognitive functions and are key pathophysiological substrates in numerous neurological and psychiatric disorders, such as Alzheimer’s disease, Parkinson’s disease, stroke, addiction and anxiety disorders. To extend this proof-of-principle work, further investigations are required to evaluate underlying mechanisms, develop strategies for improving behavioral effects and establish pathways for personalized applications with the aim of translating this exciting, innovative approach to clinical settings.

## Methods

### Participants

Forty-five healthy participants were included in the two experiments.

In Experiment 1, 15 healthy young subjects (9 females, mean±SD age 23.46±3.66 years) were recruited. Fourteen out of 15 participants performed the full protocol, while one participant dropped out between sessions for personal reasons. Only the 14 full datasets were included in the analyses.

In Experiment 2, 15 healthy older subjects (9 females, average age 66.00±4.61 years) and 15 healthy young subjects (right-handed, 8 females and 7 males, average age 26.67±4.27 years) were recruited and completed the study.

All subjects self-reported being right-handed, and handedness was confirmed by the Edinburgh Handedness Inventory^72^ (please see Table S3 in the supplementary information). The exclusion criteria are listed in the supplementary information section *“Study participant selection criteria”*. The studies were conducted in accordance with the Declaration of Helsinki. All studies were approved by the Cantonal Ethics Committee Vaud, Switzerland (*project number 2020-00127*). All participants provided written informed consent.

### Experimental protocol

Experiments 1 and 2 both followed a randomized, double-blind, crossover design. A baseline visit was always performed after inclusion, as described in the supplementary information section *“Baseline characterization of participants”* and Table S3.

### Motor task

The motor task consisted of an established and widely used 9-digit sequential finger tapping task (SFTT)^21,22^. The subjects had to reproduce a sequence shown on a computer screen with their nondominant left hand by pressing a 4-button box, with each finger corresponding to a specific number (from 2-index to 5-little finger; please see Fig. 5a). Oral and written instructions were provided, asking the participants to perform the task “*as fast and as accurately as possible”* in the fixed 30 or 90 seconds provided for each block in Experiments 1 and 2, respectively, to avoid that the participants performed the task at the extremes of their individual speed-accuracy trade-off. A dot was displayed below the number corresponding to the digit to be pressed, and the dot moved to the next digit as soon as a key was pressed, regardless of whether the correct key was chosen. No feedback about the correctness of the responses was provided. All sequences had an equivalent Kolmogorov complexity, which was determined based on a well-established procedure^22^. The order of the applied sequences before and after the crossover was randomized and counterbalanced between subjects.

### Experiment 1

In Experiment 1, four functional MRI (fMRI) sessions were performed, with two resting-state (rs) and two task-based sessions with concomitant transcranial temporal interference stimulation (tTIS) or high-frequency control stimulation (HF control). For further details, please refer to the section below. Between sessions, we included a wash-out phase of at least 3 days (7.4±4.2 days between resting state sessions and 10.3±4.7 days between task-based fMRI sessions). During the rs-MRI sessions, functional images were acquired during three resting state sequences lasting 8 minutes each, namely, before (pre), during, and after (post) stimulation, while subjects fixated on a white cross on a black background. During the task-based fMRI sessions, the participants performed six 9 minute 30 second training blocks with an approximately 1 minute 30 second break between blocks (Fig. 5b). Each block included ten 30-second repetitions of the motor task (see Experimental protocol - Motor task) with the respective stimulation condition, alternated with 30 seconds of rest (fixation cross). All participants performed a short familiarization outside the MRI environment and a 30-second baseline measurement inside the scanner before starting the training blocks. The baseline performance was verified to ensure that at least one entirely correct sequence was performed; otherwise, the baseline was repeated, and the new block was used for analysis instead of the first block. For each of the four fMRI sessions, participants completed the Stanford Sleepiness Scale^73^ (SSS) questionnaire at the beginning of the session to confirm that the subjects started the experiments at comparable levels of sleepiness across conditions (please see Fig. S4). A visual analog scale (VAS) was employed to test subjects’ attention and fatigue before and after the MRI acquisition. After each of the two study phases (i.e., resting-state and task-based fMRI), we employed a standardized questionnaire adapted from Antal and colleagues^74^ to evaluate the sensations associated with tTIS and quantify the efficiency of the blinding.

### Experiment 2

In Experiment 2, two main training sessions were performed, with at least a 3-day wash-out period between sessions (9.5±4.0 days between sessions for the younger cohort and 9.2±3.9 days between sessions for the older cohort); please see Fig. 5c for more details. During each session, either tTIS or control stimulation was applied as participants performed seven 90-second blocks of the motor task (see Experimental protocol - Motor task) alternated with 90-second breaks. In the central block, the order of requested button presses followed a predefined pseudorandom sequence to assess sequence-independent learning effects. A 90-second baseline block was acquired before training to assess initial individual performance, and 3 additional blocks were collected immediately (post), 90 minutes (FU1) and 24 hours (FU2) after the stimulation. The baseline performance was investigated to ensure that at least one entirely correct sequence was performed; this effect was consistently achieved, and thus additional repetitions were not needed. The subjects’ attention and fatigue levels were collected by employing VAS questionnaires (see above) before the baseline measurement, after the post measurement, and before and after each follow-up, while the SSS was acquired before the baseline and follow-ups to confirm that the subjects began the experimental sessions at comparable levels of sleepiness across conditions. At the end of the second post measurement, participants were asked to complete the same sensation questionnaire as for Experiment 1.

### Transcranial temporal interference stimulation (tTIS)

#### General concept

Temporal interference stimulation is a novel brain stimulation strategy that employs two or more independent stimulation channels delivering high-frequency currents (oscillating at f1 and f1+Δf) within the kHz range, which are assumed to be inert in terms of inducing neuronal activity^6,75^. The two currents generate a modulated electric field, with the envelope oscillating at the low frequency Δf (target frequency) where the currents join or cross. The peak of the envelope amplitude can be steered toward target areas located deeper in the brain by tuning the electrode position and current ratio across stimulation channels; see Grossman and colleagues^6^ and Fig. 5d for further details. Based on these properties, temporal interference stimulation can focally target deep structures without engaging overlying tissues. In the present work, we applied transcranial temporal interference stimulation (tTIS) via surface electrodes, applying a low-intensity, subthreshold protocol respecting the currently accepted cutoffs and safety guidelines for low-intensity transcranial electric stimulation^74^.

#### Stimulators

The tTIS currents were generated by two independent DS5 isolated bipolar constant current stimulators (*Digitimer Ltd, Welwyn Garden City, UK*). The stimulation patterns were created using a custom-written MATLAB-based graphical user interface and transmitted to the current sources using a standard digital-to-analog converter (*DAQ USB-6216, National Instruments, Austin, TX, USA*).

#### Stimulation paradigms

We employed two stimulation conditions: active stimulation delivered in theta bursts (tTIS) and a high-frequency control stimulation (HF control). The control stimulation consisted of two oscillatory high-frequency currents delivered at 2 kHz without any frequency shifts, which led to a flat envelope high frequency exposure incapable of eliciting brain physiological responses, as suggested by previous work^6^.

The tTIS was delivered in an intermittent pattern designed to mimic established theta-burst stimulation rhythms, which were developed in hippocampal slice preparations^23^ and have been adopted in previous NIBS approaches^24,76^. The stimulation was applied using a novel pulsed stimulation approach that utilizes frequency modulation, changing one of the two carrier frequencies to switch between modulated and unmodulated exposure. This allows to achieve an arbitrary waveform pattern without the need to change the current amplitude, thereby ensuring that the observed stimulation effects were not biased by onset effects. During tTIS, bursts of 3 peaks at 100 Hz were repeated every 200 ms (5 Hz, i.e., the theta rhythm) for 2 seconds, alternated with 8 seconds of the HF, flat envelope control (please see Fig. 5e). The other stimulation parameters were set as follows: current intensity per stimulation channel=2 mA; pure stimulation duration=30 min (for Experiment 1) and 10 min 30 sec (for Experiment 2), with breaks within each protocol; ramp-up/ramp-down period=5 sec; electrode type: round, conductive rubber with conductive cream/paste, and electrode size=3 cm^2^.

For Experiment 1, the stimulation was applied in the MRI environment by employing a standard radio frequency filter module and MRI-compatible cables (*neuroConn GmbH, Ilmenau, Germany*). The technological, safety and noise tests, and methodological factors are reported based on the ContES Checklist^77^ in Table S4 in the supplementary information.

#### Modeling

Electromagnetic simulations were performed to identify the optimal electrode placement and current steering parameters. The simulations were performed using the MIDA head model^78^, which is a detailed anatomical head model featuring 117 distinguished tissues and regions that were derived according to multimodal image data of a healthy female volunteer. Importantly, in brain stimulation modeling, the model distinguishes different scalp layers, skull layers, gray and white matter, cerebrospinal fluid, and the dura. Circular electrodes (radius=7 mm) were placed on the skin according to the 10-10 system, and the electromagnetic exposure was determined using the ohmic-current-dominated electroquasistatic solver in Sim4Life version 5.0 (*ZMT Zurich MedTech AG, Switzerland*), which is suitable due to the dominance of ohmic currents over displacement currents and the long wavelength compared to the simulation domain. The dielectric properties were assigned according to the IT’IS Tissue Properties Database v4.0^79^. Rectilinear discretization was used, and grid convergence and solver convergence analyses were performed to ensure negligible numerical uncertainty, resulting in a grid, that contained more than 54M voxels. Dirichlet voltage boundary conditions were applied, followed by current normalization, and the electrode-head interface contact was treated as ideal. tTIS exposure was simulated for 1 mA current intensity and quantified according to the maximum modulation envelope magnitude formula proposed by Grossman *et al*. in 2017^6^. Subsequently, a sweep over 960 permutations of the four electrode locations was performed, considering symmetric montages with parallel (sagittal=729 configurations; coronal=231 configurations) or crossing current paths, and the bilateral striatum (putamen, caudate, and nucleus accumbens) exposure performance was quantified according to three metrics: (1) the target exposure strength, (2) focality ratio (the volume ratio of the target tissue above the threshold to the overall brain tissue above the threshold, which measures stimulation selectivity), and (3) activation ratio (percentage of the target volume above the threshold, which measures target coverage). The threshold was defined as the 98% volumetric iso-percentile level of the tTIS. Two configurations were noted in the resulting Pareto-optimal front: one that maximized focality and activation (Pair 1: AF3 and AF4, Pair 2: TP7 and TP8 montage; focality=30.3%, activation=28.2%, threshold=0.19 V/m) and one that accepts a reduction of these two metrics by a quarter, while increasing the target exposure strength by more than 50% (Pair 1: F3 and F4, Pair 2: TP7 and TP8; focality=23.9%, activation=22.1%, threshold=0.31 V/m). The latter montage was selected because the predicted tTIS field had a larger stimulation intensity to ensure that the target could be stimulated (please see Fig. 5f).

#### Electrode positioning and evaluation of stimulation-associated sensations

The stimulation electrode positions were defined based on the above model and determined in the framework of the EEG 10-10 system^80^. The optimal positioning leading to the best stimulation of the target structure, i.e., the bilateral striatum, included the following electrodes: F3, F4, TP7 and TP8. Their scalp locations were marked with a pen. After skin preparation (cleaned with alcohol), round conductive 3 cm^2^ rubber electrodes were placed by adding a conductive paste (*Ten20, Weaver and Company, Aurora, CO, USA or Abralyt HiCl, Easycap GmbH, Woerthsee-Etterschlag, Germany*) as an interface to the skin and held in position with tape. In Experiment 1, the electrode cables were oriented toward the top of the head to allow good positioning inside the scanner, while in Experiment 2, the electrode cables were oriented toward the bottom, and fixed on the shoulders to prevent electrode displacement. The impedances were checked and optimized until they were less than 20 kΩ^64^. Once good contact was obtained, the subjects underwent current intensity testing to be familiarized with the perceived sensations and to systematically document their reactions. The tTIS and HF control stimulation protocols were both applied for 20 seconds with increasing current amplitude per channel: 0.5 mA, 1 mA, 1.5 mA and 2 mA. The participants were asked to report any kind of sensation, and if a sensation was felt, they were asked to grade the intensity from 1 to 3 (light to strong) and to provide at least one adjective to describe the sensation (please see Table S5 in the supplementary information). After this step, in Experiment 1, the cables were replaced by MRI-compatible cables, and a bandage was added to apply pressure on the electrodes and keep them in place. In Experiment 2, an EEG cap was used to hold the electrodes in place. The electrode impedances were measured before the current intensity testing, before the training with concomitant stimulation and after the intervention.

#### Image acquisition

Structural and functional images were acquired using a 3T MAGNETOM PRISMA scanner (*Siemens, Erlangen, Germany*). The 3D MPRAGE sequence was used to obtain T1-weighted images with the following parameters: TR=2.3 s; TE=2.96 ms; flip angle=9°; number of slices=192; voxel size=1×1×1 mm; and field of view (FOV)=256 mm. Anatomical T2 images were collected with the following parameters: TR=3 s; TE=409 ms; flip angle=120°; number of slices=208; voxel size=0.8×0.8×0.8 mm; and FOV=320 mm. Echo-planar imaging (EPI) sequences were used to obtain functional images with the following parameters: TR=1.25 s; TE=32 ms; flip angle=58°; number of slices=75; voxel size=2 × 2 × 2 mm; and FOV=112 mm.

#### Image preprocessing

Functional imaging data were analyzed using Statistical Parametric Mapping 12 (*SPM12; The Wellcome Department of Cognitive Neurology, London, UK*) implemented in MATLAB R2018a (*Mathworks, Sherborn, MA*). All functional images underwent the same preprocessing, including the following steps: slice time correction, spatial realignment to the first image, normalization to the standard MNI space and smoothing with a 6 mm full-width half-maximal Gaussian kernel. T1 anatomical images were coregistered to the mean functional image and then segmented to produce the forward deformation field used to normalize the functional images, allowing bias-corrected gray and white matter images to be obtained. The framewise displacement was calculated for each run to control head movement. The nonnormalized and normalized images were visually inspected to ensure good preprocessing quality. The signal-to-noise ratio was also computed to control for possible tTIS-related artifacts.

#### Signal-to-noise ratio

To verify the image quality and presence of possible artifacts due to the concomitant stimulation, total signal-to-noise ratio (tSNR) maps were computed as the mean over the standard deviation for each voxel time series. The average values of the spherical regions of interest (ROIs, 10 mm radius) underneath the four electrodes used for tTIS and underneath the theoretical positions of four more distant electrodes were extracted (please see Fig. S5 in the supplementary information). The locations of the spheres were derived by projecting the standard MNI coordinates on the scalp^81^ toward the center of the brain. The spheres were visually inspected to ensure that the whole volume was included in the brain. A linear mixed model was then used to investigate the effects of the stimulating electrodes versus those of the nonstimulating electrodes in the tSNR maps.

### Data processing

#### Resting-state MRI

Independent component analysis (ICA)-based artifact removal was performed on the preprocessed, smoothed images using the GIFT toolbox (https://trendscenter.org/software/gift/). Twenty independent components were extracted and visually inspected to remove noise-related artifacts. Seed-based connectivity analyses were implemented at the single-subject level by extracting the average time series within the striatal mask defined in the BNA atlas and including this time series as a regressor in a general linear model with six head motion parameters (three displacement motions and three rotation motions) and normalized time series in the white matter and corticospinal fluid. Regions showing significant correlation with the seed time series were extracted, and multiple comparison corrections were applied at the cluster level by controlling the FDR.

#### Task-based fMRI

A general linear model was used to estimate the signal amplitude at the single-subject level. Six head motion parameters (three displacement motions and three rotation motions) and the normalized time series in the white matter and corticospinal fluid were included as regressors. Linear contrasts were computed to estimate activation during the motor task versus that during resting periods, and ROI-based analyses were conducted. An external radiologist manually drew striatal masks on each subject’s structural T1w image. After drawing the masks for caudate and putamen, anterior and posterior sub-parts were distinguished in respect to the location of the anterior commissure. Coregistration to the functional images and normalization to MNI space were then applied to obtain individual masks for each subject. The BOLD activity within the individual striatal masks was averaged and compared between different striatal subunits, namely, putamen versus caudate, and within the putamen, namely, anterior versus posterior putamen.

Additionally, a flexible factorial design was used to compute group-level statistics, including subject, stimulation and time as factors. Multiple comparison corrections were applied at the cluster level by controlling the FDR.

#### Motor task analysis

In Experiment 1, because of the relatively short duration of the motor task repetitions (30 seconds), motor learning was evaluated by extracting the number of correct key presses per repetition divided by the number of correct key presses during the baseline measurement^82^.

In Experiment 2, motor learning was evaluated by extracting the number of correct sequences in each block divided by the correct number of sequences performed during the baseline measurement^22^. For the longer assessment blocks in Experiment 2, sequence-based outcomes were chosen instead of keypress-based outcomes because these results more closely resemble the structure of natural skilled movements, which often require smaller elements of the movements to be performed in specific order and time sequences^83^.

In both cases, frame shifts in button pressing were considered, meaning that key presses that were performed in the correct order were considered correct even if the key presses did not match the dot indicating which digit to press next.

The baseline values were compared between the stimulation conditions and among sessions to assess comparable initial performance and control for carry-over effects in each cohort; please see Fig. S2 in the supplementary information for more details.

#### Statistical analysis of the behavioral data

Statistical analyses were performed in the R software environment for statistical computing and graphics (*R Core Team (2021). R: A language and environment for statistical computing. R Foundation for Statistical Computing, Vienna, Austria. URL https://www.R-project.org/. and version 4.0.5*). To analyze the behavioral motor learning data, we conducted linear mixed effects analyses employing the *lmer* function in the *lme4* package^84^. As fixed effects, we added blocks and stimulation conditions to the model for Experiment 1 and blocks, stimulation conditions and populations (younger and older) to the model for Experiment 2. The factor subject was taken as a random intercept. Statistical significance was determined using the *anova* function with Satterthwaite’s approximations in the *lmer Test*package^85^. To mitigate the impact of isolated influential data points on the outcome of the final model, we employed tools in the *influence.ME* package to detect and remove influential points based on the following criterion: distance>4 * mean distance^86^. For specific post hoc comparisons, we conducted pairwise comparisons by computing the estimated marginal means using the *emmeans* package^87^. Effect size measures were obtained using the *effectsize* package^88^ and are expressed as_partial eta-squared (pη^2^) values for the F tests and Cohen’s d values for pairwise comparison tests, corresponding to <0.01~micro, 0.01~small, 0.06~medium, and 0.14~large effect sizes for pη^2^ and <0.2~micro, 0.2-0.3~small, 0.5~medium, and 0.8~large effect sizes for d^89^. The level of significance was set at p<0.05. Finally, for the baseline and follow-up sessions, the normality of the data distribution was tested with the Shapiro-Wilk test, and either paired t tests or Wilcoxon rank sum tests were then used. The functions were included in the *stats* package, which is part of the above referenced R software environment.

## Acknowledgements

We acknowledge access to the MRI and Neuromodulation facilities of the Human Neuroscience Platform of the Fondation Campus Biotech Geneva and access to the Neuroimaging Facilities of the HVS, Sion. This project was supported by the Defitech Foundation (Morges, CH) to F.H., the Bertarelli Foundation - Catalyst program (Gstaad, CH) to F.H., M.W., T.P., N.G. & E.N., the Novartis Research Foundation - FreeNovation (Basel, CH) to M.W. & E.N., the Wyss Center for Bio and Neuroengineering to F.H. and the Interdisciplinary Center for Clinical Research (IZKF) at the University of Würzburg (Project number Z-3R/4) to M.W. We thank M. Durand-Ruel, A. Cadic-Melchior and L. Draaisma for assistance in the randomization and R. Martuzzi, L. Mattera and O. Reynaud for support in the fMRI data acquisition.

## Author contributions

Conceptualization: M.W., E.B., T.P., F.H.; methodology: M.W., E.B., N.G., E.N., P.D., V.B., F.H.; formal analysis: M.W., E.B., V.B., P.M., I.V.; investigation: M.W., E.B., F.W., P.M., T.P., P.V., P.M.; resources: F.H., E.N.; data curation: E.B., M.W.; writing original draft/visualization: M.W., E.B.; review & editing: all co-authors; supervision: F.H.; funding: F.H., M.W., T.P., E.N., N.G.

