## Supplementary material for "LTP-like noninvasive striatal brain stimulation enhances striatal activity and motor skill learning in humans"

**Imaging results – tTIS versus HF control stimulation contrast - supporting information**

| Set-level | | Cluster-level | | | | Peak-level | | | | | x | y | z | AAL3 |
|---|---|---|---|---|---|---|---|---|---|---|---|---|---|---|
| p | c | $p_{FWE-corr}$ | $q_{FDR-corr}$ | $k_E$ | $p_{uncorr}$ | $p_{FWE-corr}$ | $q_{FDR-corr}$ | T | $(Z_E)$ | $p_{uncorr}$ | | | | |
| **0.000** | **9** | **0.015** | **0.009** | **166** | **0.001** | **0.001** | **0.004** | **6.12** | **5.76** | **0** | **-64** | **-2** | **-4** | **Temporal_Sup_L** |
| | | | | | | 0.354 | 0.172 | 4.46 | 4.31 | 0 | -58 | 12 | -10 | Temporal_Pole_Sup_L |
| | | | | | | 0.515 | 0.183 | 4.31 | 4.17 | 0 | -62 | -10 | 12 | Rolandic_Oper_L |
| | | **0.000** | **0.000** | **439** | **0.000** | **0.018** | **0.068** | **5.3** | **5.06** | **0** | **28** | **0** | **-22** | **Amygdala_R** |
| | | | | | | 0.124 | 0.151 | 4.79 | 4.61 | 0 | 30 | 6 | -8 | Putamen_R |
| | | | | | | 0.163 | 0.151 | 4.71 | 4.54 | 0 | 32 | -2 | -6 | Putamen_R |
| | | **0.012** | **0.008** | **174** | **0.001** | **0.03** | **0.073** | **5.18** | **4.95** | **0** | **10** | **-6** | **54** | **Supp_Motor_Area_R** |
| | | | | | | 0.587 | 0.209 | 4.25 | 4.12 | 0 | -6 | 0 | 60 | Supp_Motor_Area_L |
| | | | | | | 0.896 | 0.297 | 3.95 | 3.85 | 0 | 14 | 0 | 50 | Supp_Motor_Area_R |
| | | **0.001** | **0.001** | **289** | **0.000** | **0.164** | **0.151** | **4.71** | **4.54** | **0** | **-66** | **-22** | **36** | **Location not in atlas** |
| | | | | | | 0.187 | 0.152 | 4.67 | 4.5 | 0 | -60 | -22 | 30 | Postcentral_L |
| | | | | | | 0.894 | 0.297 | 3.96 | 3.85 | 0 | -60 | -28 | 22 | SupraMarginal_L |
| | | **0.011** | **0.008** | **178** | **0.001** | **0.325** | **0.172** | **4.49** | **4.34** | **0** | **-56** | **4** | **34** | **Precentral_L** |
| | | | | | | 0.915 | 0.301 | 3.93 | 3.82 | 0 | -46 | 0 | 40 | Precentral_L |
| | | | | | | 0.999 | 0.558 | 3.56 | 3.48 | 0 | -60 | 2 | 24 | Precentral_L |
| | | **0.009** | **0.008** | **184** | **0.000** | **0.354** | **0.172** | **4.46** | **4.31** | **0** | **10** | **-18** | **4** | **Thal_IL_R** |
| | | | | | | 0.469 | 0.172 | 4.35 | 4.21 | 0 | 14 | -28 | 0 | Thal_PuA_R |
| | | | | | | 0.977 | 0.376 | 3.78 | 3.69 | 0 | 18 | -20 | 8 | Thal_VPL_R |
| | | **0.021** | **0.011** | **155** | **0.001** | **0.457** | **0.172** | **4.36** | **4.22** | **0** | **-42** | **-2** | **8** | **Insula_L** |
| | | | | | | 0.937 | 0.317 | 3.89 | 3.78 | 0 | -50 | 0 | 4 | Rolandic_Oper_L |
| | | | | | | 0.999 | 0.56 | 3.55 | 3.47 | 0 | -36 | 6 | -10 | Insula_L |
| | | **0.004** | **0.006** | **215** | **0.000** | **0.563** | **0.203** | **4.27** | **4.14** | **0** | **62** | **-10** | **12** | **Rolandic_Oper_R** |
| | | | | | | 0.76 | 0.262 | 4.1 | 3.98 | 0 | 54 | -18 | 18 | Rolandic_Oper_R |
| | | | | | | 0.845 | 0.279 | 4.02 | 3.9 | 0 | 62 | -22 | 8 | Temporal_Sup_R |
| | | **0.005** | **0.006** | **207** | **0.000** | **0.641** | **0.223** | **4.21** | **4.08** | **0** | **-18** | **-50** | **-28** | **Location not in atlas** |
| | | | | | | 0.691 | 0.233 | 4.16 | 4.04 | 0 | -30 | -52 | -26 | Cerebellum_6_L |
| | | | | | | 0.957 | 0.334 | 3.84 | 3.74 | 0 | -10 | -50 | -22 | Cerebellum_4_5_L |

**Tab. S1: Summary of clusters showing higher BOLD activation for the tTIS vs. HF control contrast during the task-related fMRI experiment (Experiment 1)**

**BOLD activity in control regions beneath the electrodes**

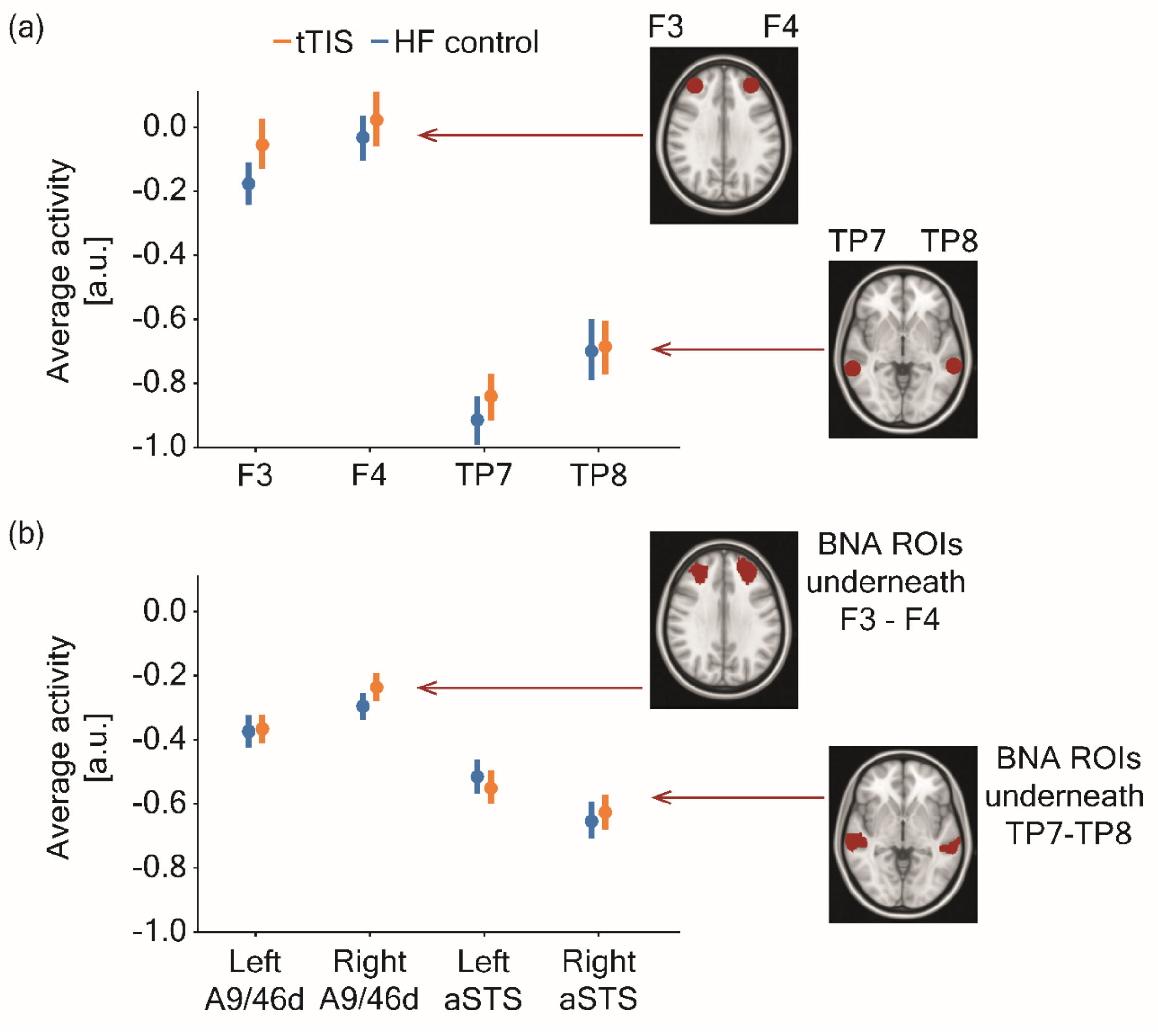


**Fig. S1: BOLD activity in control regions beneath the electrodes**
BOLD activity during the task-based fMRI experiment (Experiment 1) was extracted and averaged across blocks from **(a)** spheres (radius=10 mm) placed below the stimulation electrodes (EEG 10-10 positions: F3, F4, TP7, TP8) or **(b)** corresponding regions based on the Brainnetome (BNA) Atlas[1] (the left and right dorsal area A9/46d for the F3/F4 or the left and right anterior superior temporal sulcus (aSTS) for the TP7/TP8 positions). Please note that the values are negative as the activity during the task is referenced to activity during rest periods, in which the subjects kept their eyes open and fixated a cross, following a standard block design procedure. Spheres model: $F(1,651)=2.04$, $p=0.15$; BNA model $F(1,651)=0.38$, $p=0.54$.

**Imaging results – parametric modulation - supporting information**

| Set-level | | Cluster-level | | | | Peak-level | | | | | x | y | z | AAL3 |
|---|---|---|---|---|---|---|---|---|---|---|---|---|---|---|
| p | c | $p_{FWE\text{-}corr}$ | $q_{FDR\text{-}corr}$ | $k_E$ | $p_{uncorr}$ | $p_{FWE\text{-}corr}$ | $q_{FDR\text{-}corr}$ | T | $(Z_E)$ | $p_{uncorr}$ | | | | |
| **0.000** | | **0.000** | **0.004** | **197** | **0.000** | **0.133** | **0.076** | **8.35** | **4.83** | **0** | **28** | **4** | **-4** | **Putamen_R** |
| | | | | | | 0.910 | 0.366 | 6.06 | 4.11 | 0 | 26 | 16 | 2 | Putamen_R |
| | | | | | | 0.985 | 0.530 | 5.58 | 3.92 | 0 | 28 | 4 | 10 | Putamen_R |

**Tab. S2: Summary of clusters showing stronger modulation of BOLD activity by behavioral performances within the right striatum**

**Motor performance at baseline or during pseudorandom sequences**

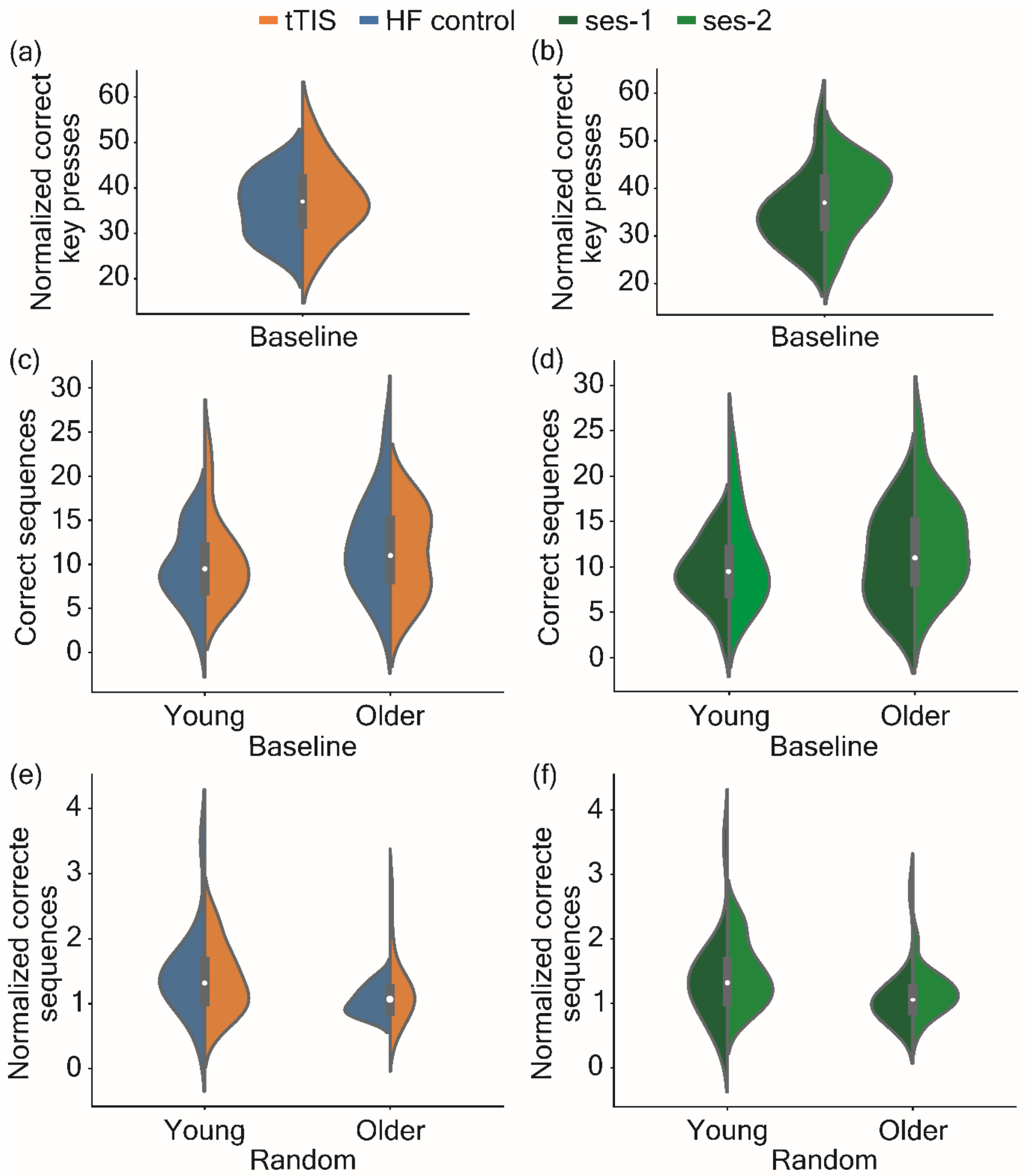


**Fig. S2: Motor performance at baseline or during pseudorandom sequences**
**(a)** Comparison of baseline performance between tTIS and HF control stimulation in Experiment 1. No significant difference was found (*t(13)=0.29, p=0.78, d=0.08; Bayesian paired t-test,* $BF_{10}$*=0.28* [*moderate evidence in favor of the Null Hypothesis (*$H_0$*)*]). **(b)** Comparison of baseline performance between first and second training session in Experiment 1. A significant difference was found, with higher performance during the second session (*t(13)=-3.35, p=0.005, d=-0.77; Bayesian paired t-test,* $BF_{10}$*=9.769* [*moderate evidence in favor of the Alternative Hypothesis (*$H_1$*)*]). **(c)** Comparison of baseline performance between tTIS and HF control stimulation in Experiment 2 for the young cohort on the left and the older cohort on the right. No significant difference was found (young: *t(14)=0.55, p=0.59, d=0.22; Bayesian paired t-test, BF10=0.373* [*anecdotal evidence for* $H_0$]; older: *t(14)=-0.55, p=0.59, d=-0.21; Bayesian paired t-test, BF10=0.368* [*anecdotal evidence for* $H_0$]). **(d)** Comparison of baseline performance between first and second training session in Experiment 2 for the young cohort on the left and older cohort on the right. No significant difference was found in the young cohort

(*t(14)=-1.44, p=0.17, d=-0.34; Bayesian paired t-test, BF10=0.620* [*anecdotal evidence for* $H_0$]), whilst a marginal significant increase in performance was observed in the older cohort during the second session (*t(14)=-2.10, p=0.05, d=-0.45; Bayesian paired t-test, BF10=1.439* [*anecdotal evidence for* $H_1$]). **(e)** Comparison of performance during a pseudorandom motor sequence between tTIS and HF control stimulation in Experiment 2 for the young cohort on the left and the older cohort on the right. No significant difference was found (young: *V=53, p=0.72, d=-0.19; Bayesian paired t-test, BF10=0.29* [*moderate evidence for* $H_0$]*;* older: *V=82, p=0.23, d=0.44; Bayesian paired t-test, BF10=0.56* [*anecdotal evidence for* $H_0$]). **(f)** Comparison of performance during a pseudorandom motor sequence between the first and the second training session in Experiment 2 for the young cohort on the left and the older cohort on the right. No significant difference was found (young: *V=50, p=0.60, d=-0.10; Bayesian paired t-test, BF10=0.269* [*moderate evidence for* $H_0$]*;* older: *V=32, p=0.21, d=-0.17; Bayesian paired t-test, BF10=0.292* [*moderate evidence for* $H_0$]).

**Evaluation of tTIS-associated sensations**

Stimulation-associated sensations of tTIS and HF control stimulation were systematically studied and characterized as they potentially impose a challenge for the blinding integrity of controlled neuromodulation studies and might lead to bias[2,3]. The analysis of the reported sensations during the stimulation tests preceding the main experiment (N=119) indicated a comparable level of stimulation-associated sensations across the tested stimulation conditions (tTIS vs. HF control) at the tested intensity levels (0.5 to 2 mA per stimulation channel). This was indicated by a non-significant stimulation condition x current strength interaction ($F(3,837.95)=0.06$, $p=0.98$, $p\eta^2<0.001$ [micro], *BF10=0.001,* [*decisive evidence for* $H_0$]), please see also Fig. S3a&b. The subjective characterization of the quality of the perceived sensations, as reported by the subjects, are listed in Tab. S5 below. Furthermore, the post-interventional assessments of TI-associated sensations perceived by the subjects during the intervention phases (fMRI plus stimulation or motor training plus stimulation) revealed a similar frequency of the evaluated sensation categories across the tested stimulation conditions (stimulation condition x sensation category $F(6,727.26)=0.73$, $p=0.63$, $p\eta^2=0.006$ [micro], *BF10=0.0009* [*decisive evidence for* $H_0$]), please see Fig. S3c&d. In an additional step, we systematically asked the subjects to provide their best guess on the applied stimulation condition. The analysis of the tTIS sessions suggested that in both, Experiment 1 and Experiment 2, the frequency of correct and incorrect responses was not significantly different from each other (exact binomial test for (i) rs-fMRI: *p=0.55*, (ii) task-based fMRI: *p=0.75*, (iii) behavioral experiment young subjects: *p=1.00*, (iv) behavioral experiment older subjects: *p=1.00*). In conclusion, the analyses showed an excellent blinding integrity of the novel tTIS protocol with respect to the HF control condition.

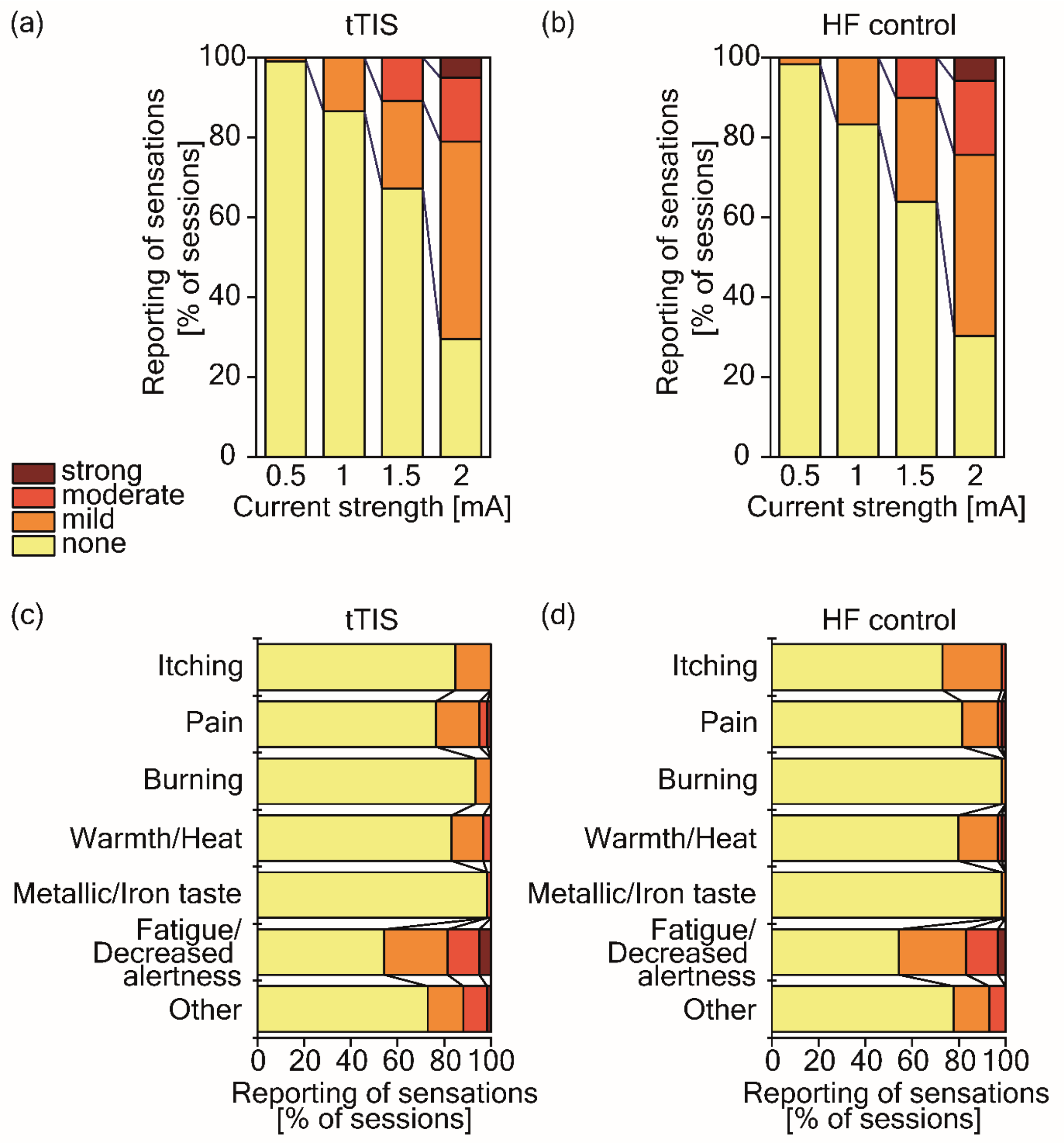


**Fig. S3: tTIS-associated sensations**

(**a&b**) Depicted is the frequency of reported sensations separated by their strength (none, mild, moderate, strong) for the tested currents intensity levels (0.5, 1.0, 1.5, 2 mA per stimulation channel) during the stimulation-associated sensation tests preceding a total of N=119 experimental sessions. No differences in perceived stimulation-associated sensations could be detected comparing the two tested conditions tTIS and HF control, as indicated by a non-significant stimulation condition x current strength interaction ($F(3,837.95)=0.06$, $p=0.98$, $p\eta^2<0.001$ [micro]). (**c&d**) Systematic evaluation of the perceived sensations during the interventional sessions (N=118) revealed no stimulation-condition associated differences for the tested commonly with tES-protocols associated sensations[4] (stimulation condition x sensation category $F(6,727.26)=0.73$, $p=0.63$, $p\eta^2=0.006$ [micro]).

**Study participant selection criteria**

**Exclusion criteria**

- Unable to consent
- Severe neuropsychiatric (e.g., major depression, severe dementia) or unstable systemic diseases (e.g., severe progressive and unstable cancer, life threatening infectious diseases)
- Severe sensory or cognitive impairment or musculoskeletal dysfunctions prohibiting to understand instructions or to perform the experimental tasks
- Inability to follow or non-compliance with the procedures of the study
- Contraindications for noninvasive brain stimulation (NIBS) or magnetic resonance imaging (MRI):
  - Electronic or ferromagnetic medical implants/device, non-MRI compatible metal implant
  - History of seizures
  - Medication that significantly interacts with NIBS being benzodiazepines, tricyclic antidepressant and antipsychotics
- Regular use of narcotic drugs
- Left-handedness
- Pregnancy
- Request of not being informed in case of incidental findings
- Concomitant participation in another trial involving probing of neuronal plasticity.

**Baseline characterization of participants**

Baseline questionnaires:

- Demographic data (age, sex),
- Handedness (Edinburgh Laterality Test)[6],
- Sleep quality (Pittsburg Sleep Quality Index, PSQI)[7],
- Cognition (Frontal Assessment Battery, FAB)[8] - for older cohort only,
- Cognition (MOntreal Cognitive Assessment, MOCA)[9] - for older cohort only.

| Experiment | Age | Sex | Handedness | PSQI | FAB | MOCA |
|---|---|---|---|---|---|---|
| #1 | 23.46 ± 3.66 | 8/15 females | 87.68 ± 16.59 | 4.50 ±1.79 | n/a | n/a |
| #2 (older subjects) | 66.00 ± 4.61 | 9/15 females | 89.00 ± 14.47 | 3.47 ± 1.81 | 16.20 ± 2.11 | 27.67 ± 1.76 |
| # 2 (young subjects) | 26.67 ± 4.27 | 9/15 females | 79.26 ± 23.82 | 5.67 ± 1.95 | n/a | n/a |

**Tab. S3: Overview of participants' characteristics**

**Questionnaires on attention, fatigue and sleepiness**

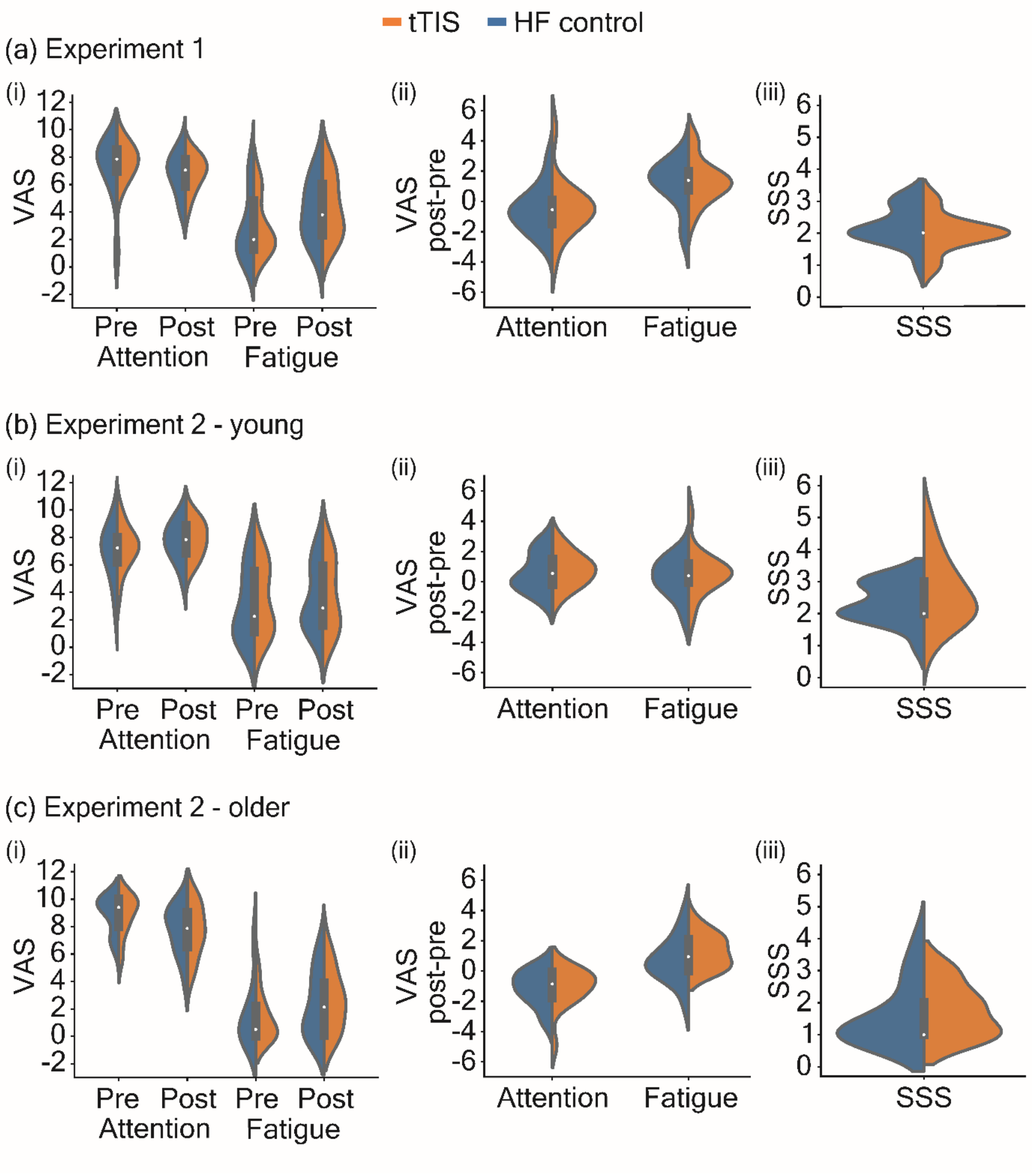


**Fig. S4: Questionnaires on attention, fatigue and sleepiness**
**(a)** Subjects' level of attention and fatigue during Experiment 1, quantified with a visual analogue scale (VAS) ranging from 0 to 10. (i) Comparison of attention and fatigue levels between tTIS and HF control stimulation sessions, pre and post training. No significant difference was found (attention pre: $V=54, p=0.95$; attention post: $t(13)=-0.06, p=0.95$; fatigue pre: $V=50.5, p=0.93$; fatigue post: $t(13)=-0.24, p=0.82$). (ii) Comparison of changes in attention and fatigue levels pre-post training between tTIS and HF control stimulation sessions. No significant difference was found (attention: $V=45, p=0.66$; fatigue: $t(13)= -0.77, p=0.46$). (iii) Comparison of sleepiness, quantified with the Stanford Sleepiness Scale (SSS)[5], between tTIS and HF control stimulation sessions, pre training. No significant difference was found ($V=24, p=0.41$). **(b)** Attention and fatigue level in Experiment 2 in the young population, the same assessment method as in experiment 1 was used. (i) Comparison of attention and fatigue levels between tTIS and HF control stimulation sessions, pre and post training. No significant difference was found (attention pre: $t(14)=0.54, p=0.60$; attention post: $t(14)=0.36, p=0.73$; fatigue pre: $V=68.5,$

*p=0.65*; fatigue post: V=*63, p=0.89*). (ii) Comparison of changes in attention and fatigue levels pre-post training between tTIS and HF control stimulation sessions. No significant difference was found (attention: *t(14)=-0.54, p=0.60*; fatigue: *t(14)=-1.02, p=0.32*). (iii) Comparison of sleepiness between tTIS and HF control stimulation sessions, pre training. No significant difference was found (V=16, p=0.24). **(c)** Attention and fatigue level in Experiment 2 in the older population. (i) Comparison of attention and fatigue levels between tTIS and HF control stimulation sessions, pre and post training. No significant difference was found (attention pre: *V=53, p=1*; attention post: *t(14)=0.06, p=0.95*; fatigue pre: *V=65.5, p=0.78*; fatigue post: *V=56, p=0.85*). (ii) Comparison of changes in attention and fatigue levels pre-post training between tTIS and HF control stimulation sessions. No significant difference was found (attention: *V=35, p=0.48*; fatigue: *t(14)=-1.55, p=0.14*). (iii) Comparison of sleepiness between tTIS and HF control stimulation sessions, pre training. No significant difference was found (V=22.5, p=0.64).

**ContES Checklist**

| **Technological factors** | |
|---|---|
| Manufacturer of Stimulator | DS5 Isolated Bipolar Constant Current Stimulator *(Digitimer Ltd, Welwyn Garden City, UK)* |
| MR Conditional Electrodes | Round, 3 $cm^2$ conductive rubber electrodes |
| Electrode Positioning | F3>F4<br>TP7>TP8<br><br>A bandage is warped around the head to apply pressure and keep the electrodes in place<br><br>Electrodes are oriented in order to have vertical cables entering parallel to the MRI coil<br><br>Head was fixed with pillows to avoid movements |
| MR Conditional Skin-Electrode Interface | Ten20 conductive paste *(Weaver and Company, Aurora, CO, USA)*<br><br>One or two drops of saline were added when impedances were too high |
| Amount of Contact Medium (Paste/Gel/Electrolyte) | Around 1 mm of paste was manually placed on the electrodes |
| Electrode Placement Visualization | Pictures<br>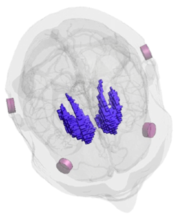 |
| RF Filter | NeuroConn DC-STIMULATOR MR RF filter module with MRI-compatible cables and electrodes *(neuroConn GmbH, Ilmenau, Germany)* |
| Wire Routing Pattern | 10 m ethernet cables between inner and outer box pass through a conduit along the wall of the MRI room until reaching the back of the MRI. Cables are then fixed with straps on the ground and on the wall of the MRI machine in order to avoid loops until reaching the interior of the coil.<br><br>Cables between the head and the inner boxes were also fixed with straps and they were oriented in |

| | |
|---|---|
| | order to minimize the magnetic field influence as soon as possible, as indicated by the red arrows of the image below.<br>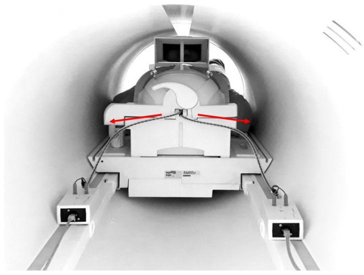 |
| tES-fMRI Machine Synchronization/Communication | Stimulation was triggered by the stimulus delivery PC via parallel port to BNC cable. The parallel port of the stimulus delivery PC was connected to the DAQ controlling the stimulators. Stimulus delivery PC, in turn, also received the scanner trigger from the scanner via USB port. |
| **Safety and noise tests** | |
| MR Conditionality Specifics for tES Setting | Please refer to Section *"Methods: Imaging acquisition"* |
| tES-fMRI Setting Test - Safety Testing | Impedances were checked before and after the stimulation.<br><br>No temperature tests were performed during the experiment.<br><br>Intensity titration was performed prior to entering the MRI, testing increasing currents (0.5, 1, 1.5 and 2 mA) and asking the subject to report any type of sensation.<br><br>A sensation questionnaire was also performed at the end of the experiment. |
| tES-fMRI Setting Test - Subjective Intolerance Reporting | No intolerances were reported by any subject |
| tES-fMRI Setting Test - Noise/Artifact | Signal to Noise Ratio (SNR) analyses were performed on the fMRI images, please refer to Section *"Preprocessing quality control"* below. |
| Impedance Testing | Impedances were checked right after electrode positioning outside the scanner, before and after the stimulation inside.<br><br>One or two drops of saline solution were added if impedances were higher than 20 kΩ |

| Methodological factors | |
|---|---|
| Concurrent tES-fMRI Timing | For timings, please refer to Fig. 1b in main text of the manuscript |
| Imaging Session Timing | All sequences were performed with tTIS electrodes placed on the subjects' head. |
| tES Experience Report | Please refer to Section *"Evaluation of tTIS-associated sensations" above.* |

**Tab. S4: ContES Checklist**
Reporting on technological, safety and noise tests, and methodological factors based on the ContES Checklist.[10]

**Evaluation of stimulation-associated sensations**

| Experiment | HF control | tTIS |
|---|---|---|
| #1 | • Tingling | • Tingling |
| | • Warm/Heat | • Warm |
| | • Burning | |
| | • Shiver | • Shiver |
| | • Vibration | • Vibration |
| | • Pain | • Pain |
| | • Waves | • Waves |
| | • Itching | • Itching |
| | • Scratching | • Scratching |
| | • Pressure | • Pressure |
| | • Massaging | • Massaging |
| | • Tickling | • Tickling |
| | • Tiny needles | • Tiny needles |
| | | • Slight touch |
| | | • Stinging |
| #2 | *Young subjects:* | *Young subjects:* |
| | • Tingling | • Tingling |
| | • Pressure | • Pressure |
| | • Slight touch | |
| | • Warm | |
| | • Pinching | • Pinching |
| | • Tickling | • Tickling |
| | • Ants | • Ants/like a small insect moving |
| | • Massaging | |
| | • Contracting | • Contracting |
| | • Pins and needles | |
| | • Drilling | • Drilling |
| | • Vibration | • Vibration |
| | • Pulsating | |
| | | • Shiver |
| | | • Goose bumps |
| | *Older subjects:* | *Older subjects:* |
| | • Warm | • Warm |
| | • Oscillation | • Oscillation |
| | • Tingling | • Tingling |
| | • Pain | |
| | • Tickling | • Tickling |
| | • Pressure | • Pressure |

**Tab. S5: tTIS associated sensations – subjective quality based on subjects' reports**

**Preprocessing quality control**

A threshold of 0.5 was chosen to discard subjects showing more than 40% of voxels with framewise displacement (FD) higher than this threshold. In the current study cohort, no subject exceeded the limit value, thus the whole dataset could be used. Furthermore, successful cleaning of the data were ensured by visual checking the preprocessing results. In particular, good registration between anatomical and functional images and normalization to standard space were checked.
The signal to noise ratio analysis showed significantly higher tSNR values underneath the stimulating electrodes ($F(1,1323)=564.68$, $p<0.001$ , $p\eta^2=0.3$ [large]), see Fig. S5. This result indicates that the stimulation did not introduce additional noise in the MR images, since this would have led to lower SNR values. In summary, all controls confirmed the good quality of the imaging data.

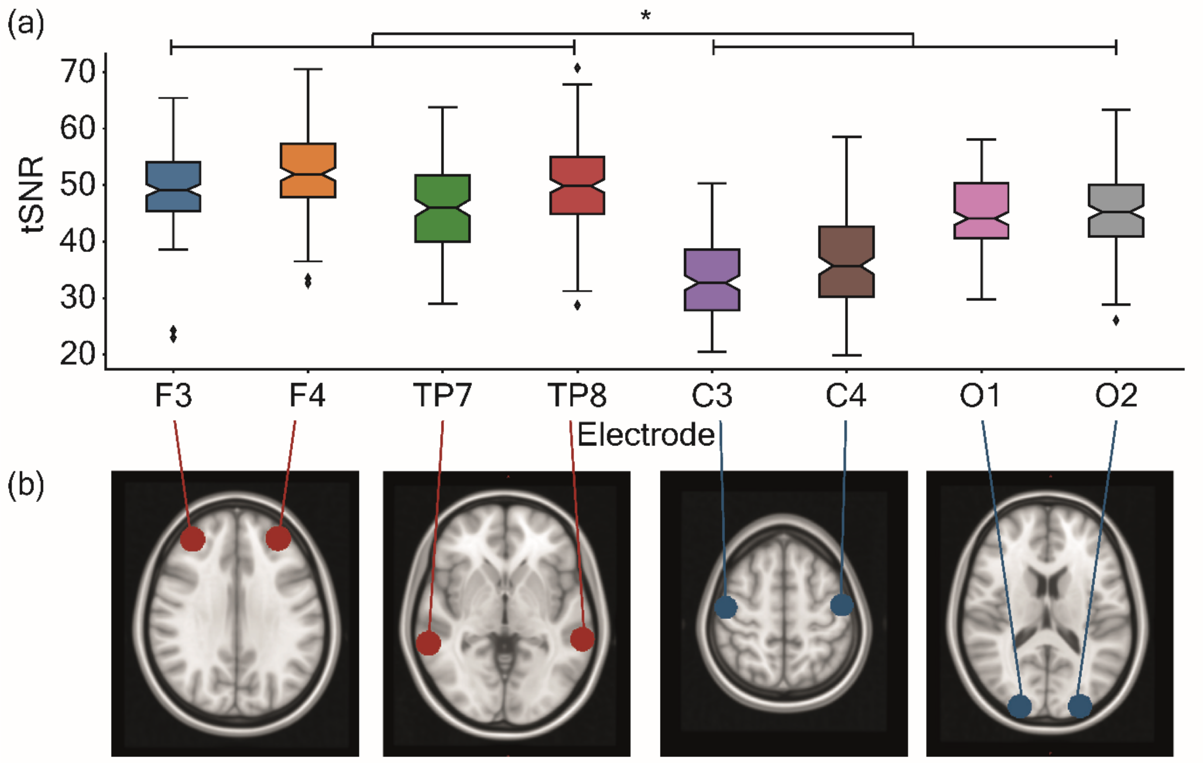


**Fig. S5: Signal to noise ratio analysis**
Total signal to noise ratio (tSNR) maps were created in order to investigate possible stimulation-induced artifacts. tSNR values were obtained as the ratio between the mean and the standard deviation of the time series during each fMRI block for each subject. An average value was then extracted from spheres of 10 mm radius underneath each of the four stimulation electrodes and underneath other four control electrode positions (not used in the study). **(a)** tSNR average values are plotted for each of the electrode-associated sphere. Stimulation electrodes showed higher tSNR with respect to the control electrodes *($F(1,1323)=564.68$, $p<0.001$ , $p\eta^2=0.3$ [large])*, indicating no additional noise was introduced by the stimulation. **(b)** Images showing the position of the chosen spheres, underneath the used electrodes (in red) and underneath the control positions (in blue).
